# Organelle proteomic profiling reveals lysosomal heterogeneity in association with longevity

**DOI:** 10.1101/2022.10.16.512400

**Authors:** Yong Yu, Shihong M. Gao, Youchen Guan, Pei-Wen Hu, Qinghao Zhang, Jiaming Liu, Bentian Jing, Qian Zhao, David M Sabatini, Monther Abu-Remaileh, Sung Yun Jung, Meng C. Wang

**Author notes:** Equal contribution (Yong Yu, Shihong M. Gao, Youchen Guan). Correspondence and requests for materials should be addressed to Meng C. Wang or Yong Yu.

## Abstract

Lysosomes are active sites to integrate cellular metabolism and signal transduction. A collection of proteins associated with the lysosome mediate these metabolic and signaling functions. Both lysosomal metabolism and lysosomal signaling have been linked to longevity regulation; however, how lysosomes adjust their protein composition to accommodate this regulation remains unclear. Using deep proteomic profiling, we systemically profiled lysosome-associated proteins linked with different longevity mechanisms. We further discovered the lysosomal recruitment of AMPK and nucleoporin proteins and their requirements for longevity in response to increased lysosomal lipolysis. Through comparative proteomic analyses of lysosomes from different tissues and labeled with different markers, we discovered lysosomal heterogeneity across tissues as well as the specific enrichment of the Ragulator complex on Cystinosin positive lysosomes. Together, this work uncovers lysosomal proteome heterogeneity at different levels and provides resources for understanding the contribution of lysosomal protein dynamics to signal transduction, organelle crosstalk and organism longevity.

## INTRODUCTION

Lysosomes are membrane-bound organelles that are specialized to constitute an acidic environment in the cytosol. Lysosomes carry many of proteins that are essential for maintaining lysosomal activities and mediating lysosomal regulatory effects. Inside the lysosomal lumen, a series of acidic hydrolases, including lipases, proteases, glucosidase, acid phosphatase, nuclease and sulfatases, are responsible for the degradation and recycling of extracellular and intracellular materials delivered through endocytic, phagocytotic and autophagic processes (Appelqvist et al., 2013; Ballabio & Bonifacino, 2020; Lawrence & Zoncu, 2019). Additionally, on the lysosomal limiting membrane, a group of integral transmembrane proteins play crucial roles in the maintenance of luminal acidic pH and ion homeostasis, the control of lysosomal membrane potential and export of metabolic products, as well as the regulation of organelle interaction and signal transduction (Ballabio & Bonifacino, 2020; Lawrence & Zoncu, 2019). For example, the lysosomal vacuolar-type H^+^-ATPase (v-ATPase) on the membrane is the primary driver for the active accumulation of protons in the lysosomal lumen, which also requires a neutralizing ion movement mediated by ion channels and transporters (Graves et al., 2008; Nicoli et al., 2019). In addition, v-ATPase coordinates with lysosomal amino acid transporter SLC38A9 and lysosomal cholesterol exporter NPC1 in regulating the activation of mechanistic/mammalian target of rapamycin complex I (mTORC1) by amino acid and lipid cues (Castellano et al., 2017; Wang et al., 2015). The recruitment of mTORC1 to the lysosome is mediated by RagA/B and RagC/D GTPase heterodimers that are associated with the scaffold protein complex Ragulator tethered on the lysosomal membrane (de Araujo et al., 2017). Through interacting with Axin, Ragulator also mediates the activation of AMP-activated protein kinase (AMPK) on the lysosomal surface (Zhang et al., 2014). Furthermore, lysosomes are not static, isolated organelles; instead they are highly mobile vesicles that undergo frequent movements in both anterograde (nucleus-to-periphery) and retrograde (periphery-to-nucleus) directions and form dynamic interactions with other organelles including endosomes, autophagosomes, endoplasmic reticulum and mitochondria (Ballabio & Bonifacino, 2020; Pu et al., 2016). These trafficking and interaction processes are mediated by lysosomal integral transmembrane proteins as well as diverse proteins that are recruited to lysosomes in response to different extracellular and intracellular inputs (Ballabio & Bonifacino, 2020; Pu et al., 2016).

Lysosomes control numerous cellular processes, and dysfunction of lysosomes has been linked with various diseases, such as lysosomal storage disorders (Ballabio & Gieselmann, 2009; Platt et al., 2012), Alzheimer’s disease (Nixon & Cataldo, 2006), Parkinson disease (Navarro-Romero et al., 2020) and some types of cancer (Davidson & Vander Heiden, 2017; Fehrenbacher & Jaattela, 2005). Emerging evidence also suggests the lysosome as a central regulator of organism longevity, through its involvement in autophagy and its modulation of metabolic signaling pathways. The induction of autophagic flux has been observed in multiple pro-longevity states, and is required for the pro-longevity effects caused by those genetic, dietary and pharmacological interventions, such as reduced insulin/IGF-1 signaling, caloric restriction and spermidine treatment (Hansen et al., 2018). On the other hand, lysosomes are now recognized as the key platform to modulate the activities of mTORC1 and AMPK signaling, two well-characterized longevity regulating pathways (Savini et al., 2019). In addition, our studies have discovered lysosomal lipid messenger pathways that are induced by a lysosomal acid lipase LIPL-4 and promote longevity via both cell-autonomous and cell-nonautonomous signaling mechanisms (Folick et al., 2015; Ramachandran et al., 2019; Savini et al., 2022; Wang et al., 2008). Given the importance of lysosomes in regulating longevity, it will be crucial to understand how changes in the lysosomal protein composition are associated with longevity regulation.

To systemically profile the protein composition of lysosomes, methods have been developed to purify lysosomes using gradient centrifugation (Gao et al., 2017; Lubke et al., 2009; Markmann et al., 2017; Schroder et al., 2010). More recently, a lysosome immunoprecipitation method, which uses anti-HA (human influenza virus hemagglutinin) antibody conjugated magnetic beads to immune-purify lysosomes from mammalian cells expressing transmembrane protein 192 (TMEM192) fused with three tandem HA (3×HA) epitopes, has further improved the specificity and speed of lysosomal isolation (Abu-Remaileh et al., 2017). This rapid isolation method has facilitated follow-up mass spectrometry (MS)-based proteomics as well as metabolomics analyses (Abu-Remaileh et al., 2017; Eapen et al., 2021; Laqtom et al., 2022).

In the present study, we have applied an immunoprecipitation-based method for rapid isolation of lysosomes from live *C. elegans* using transgenic strains expressing lysosomal membrane proteins tagged with 3×HA (Lyso-Tag). We then conducted large-scale proteomic profiling using isolated lysosomes and remaining non-lysosomal fractions, to determine the enrichment of each identified protein on the lysosome. Based on these analyses, we have defined a lysosome-enriched proteome and compared it between wild-type and long-lived worms, revealing lysosomal protein composition changes associated with longevity. We have also generated transgenic strains expressing Lyso-Tag specifically in the hypodermis (epidermal), muscle, intestine (fat storage/digestion) and neurons, leading to the discovery of lysosomal proteome heterogeneity in different tissues. Furthermore, by comparing the lysosome-enriched proteome with LAMP1/LMP-1 Lyso-Tag and the one with Cystinosin/CTNS-1 Lyso-Tag, we discovered that the Ragulator complex and other mTORC1 regulators exhibit increased enrichments on lysosomes containing the cysteine transporter Cystinosin.

## RESULTS

### Map lysosome-enriched proteome systemically in *C. elegans*

To comprehensively reveal proteins that are enriched at the lysosome, we have applied rapid lysosome immunoprecipitation followed by MS-based proteomic profiling (Lyso-IP) (Figure 1A). We first generated a transgenic strain expressing the lysosome-associated membrane protein, LMP-1 (Eskelinen, 2006) fused to both 3×HA and RFP (LMP-1 *LysoTg*). Fluorescence imaging of RFP confirmed the lysosomal localization of the LMP-1 fusion protein in live organisms and made it possible to follow purified lysosomes *in vitro* (Figure 1B). The 3×HA epitope tag is used to purify lysosomes from homogenized worm lysate via immunoprecipitation using anti-HA antibody conjugated magnetic beads (Figure 1A). In general, about 160,000 worms at day-1 adulthood were harvested and homogenized. Upon centrifugation to remove debris and nuclei, 3×HA-tagged lysosomes were immunoprecipitated and separated from other cellular content (flow-through controls, Figure 1A). The whole process from harvesting worms to purified lysosomes takes around 25 minutes. Many purified lysosomes were able to take up LysoTracker probes and exhibit positive fluorescence signals, indicating that they maintain intact with an acidic pH; while there are also some broken lysosomes losing LysoTracker staining (Figure 1C). When blotting with antibodies against different organelle markers, we found that the purified lysosomes show no or nearly no protein markers of other organelles, including HSP-60 (mitochondria heat shock protein) (Hartl et al., 1992; Mayer, 2010), CYP-33E1 (Endoplasmic Reticulum (ER) cytochrome P450) (Brown & Black, 1989), SQV-8 (Golgi glucuronosyltransferase) (Hadwiger et al., 2010) and beta-actin (cytoskeleton) (Figure 1D), while the flow-through controls show these protein markers but nearly no lysosomal protein marker LMP-1 (Figure 1D). Together, these results demonstrate the efficacy of the Lyso-IP approach to enrich for lysosomal proteins.

**Figure 1.**
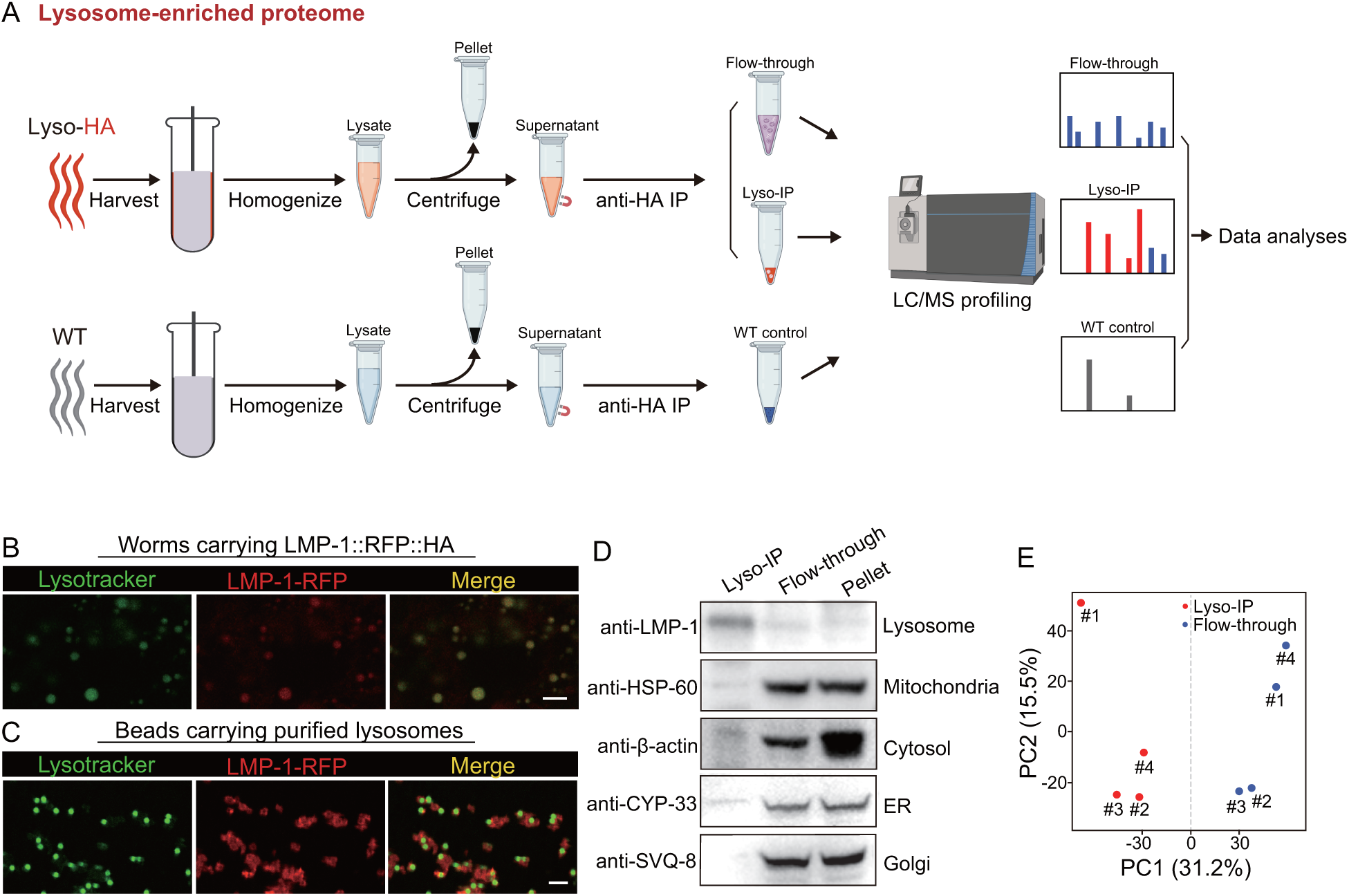
Rapid lysosome isolation coupled with proteomic profiling. (**A**) Schematic of the workflow for immunoprecipitation-based lysosome purification (Lyso-IP) and mass-spectrometry based proteomic profiling to identify lysosome-enriched proteomes in *C. elegans*. (B) Example images of transgenic strains carrying LMP-1 Lyso-Tag (LMP-1::RFP-3×HA) with LysoTracker staining to mark lysosomes *in vivo*. Scale bar=5 µm. (C) Example images of beads carrying purified lysosomes from Lyso-IP with LysoTracker staining to mark intact lysosomes *in vitro.* Scale bar=5 µm. (D) Western blot for protein markers of different subcellular compartments using purified lysosomes (Lyso-IP), paired non-lysosomal fractions (Flow-through) or Pellet. (E) PCA analysis of four independent biological replicates of Lyso-IP and Flow-through samples.

Next, we conducted proteomic profiling of purified lysosomes with their paired flow-through controls (Figure 1A). The correlation analysis shows good reproducibility among four independent biological replicates (Supplementary Figure 1A), and the PCA analysis shows a clear separation between Lyso-IP replicates and flow-through controls (Figure 1E). In parallel, we also conducted immunoprecipitation using homogenized lysate from wild-type (WT) worms that do not carry a Lyso-Tag and then analyzed proteomic profiles of three independent samples as non-tag controls (Figure 1A). Based on these proteomic data, we used three criteria to identify lysosome-enriched proteins: first, their levels in the purified lysosomes are 10-fold or higher than those in the flow-through controls (Figure 2A); second, their enrichments can be repeated in all biological replicates (Figure 2A); and lastly, their enrichments over non-tag controls are more than 2-fold (Figure 2B).

**Figure 2.**
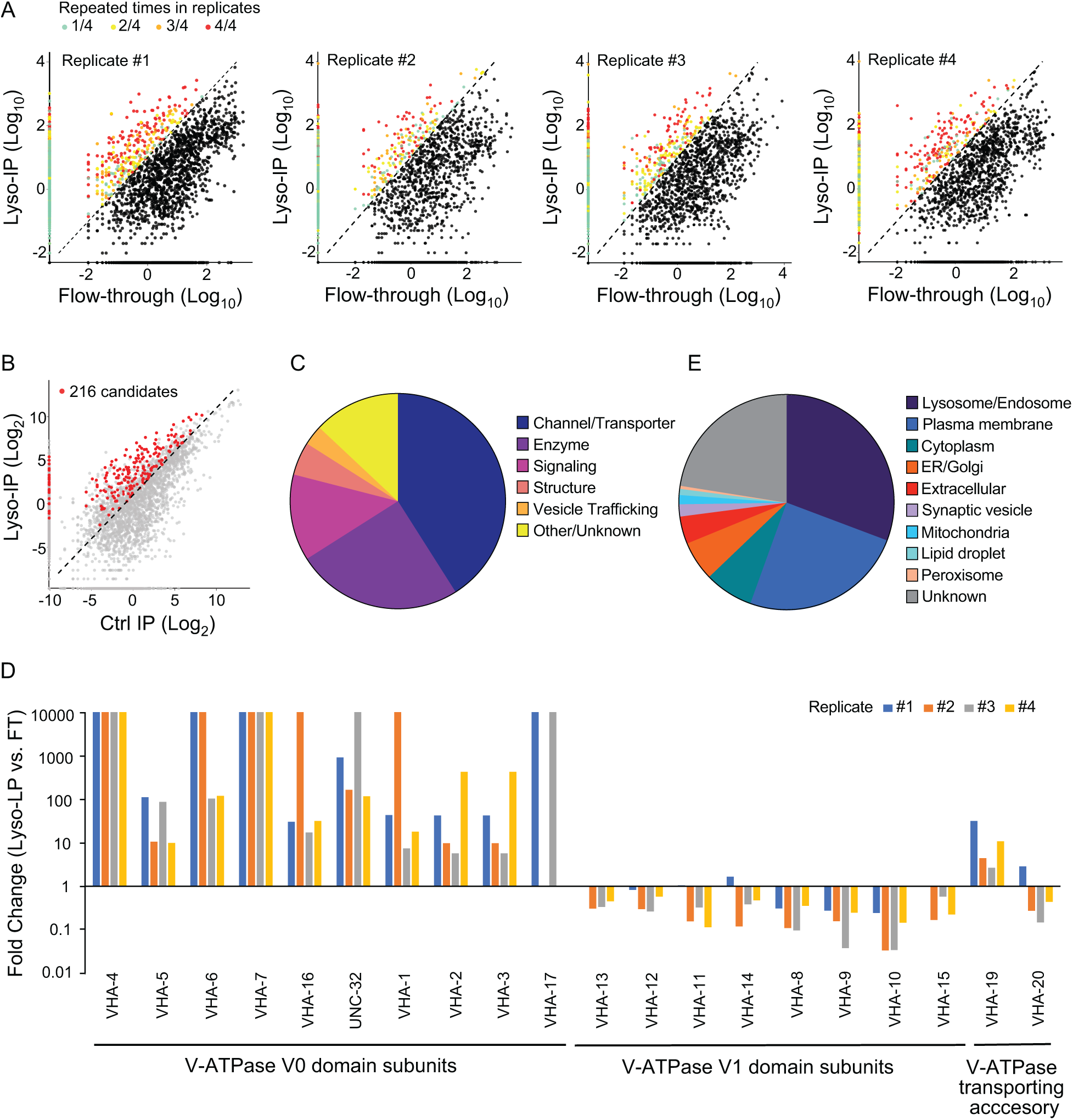
Systemic view of lysosome-enriched proteome. (**A**) Scatter plots showing candidate selection from four independent biological replicates in proteomics analyses. Proteins with at least 10-fold higher levels in Lyso-IP samples than in flow-through controls are highlighted with different colors based on repeated times in four replicates. (B) Scatter plot showing candidate selection with normalization to non-tagged controls using wild-type worms. 216 proteins with over 2-fold higher levels in Lyso-IP samples than in non-tagged controls are highlighted in red. (C) Pie chart showing molecular function categories of lysosome-enriched proteins. (D) The lysosomal enrichment ratio (Lyso-IP vs FT) for each subunit of lysosomal vacuolar ATPase (v-ATPase) in four independent replicates is shown. (E) Pie chart showing subcellular location categories of lysosome-enriched proteins.

Together, 216 lysosome-enriched candidates were identified from more than 6000 detected proteins, and 178 candidates have mammalian homologs (Supplementary Figure 1B, Supplementary Table 1). This lysosome-enriched proteome consists of 83 membrane transporters and channels, 47 enzymes, 26 signaling factors, 12 structural components, and 6 involved in vesicle trafficking (Figure 2C). These include known lysosomal proteins, such as various lysosomal Cathepsins that catalyze protein degradation (Turk et al., 2012), lysosomal specific ARL8 GTPase that mediates lysosome trafficking (Nakae et al., 2010), and subunits of lysosomal v-ATPase that pumps protons into the lysosomal lumen to maintain an acidic pH (Forgac, 2007) (Supplementary Table 1).

Lysosomal v-ATPase consists of both V0 and V1 domains that are associated with the lysosomal membrane and faced the cytosol, respectively. Reversible dissociation of the V1 and V0 domains responds to nutritional signals and plays a crucial role in the regulation of the lysosomal v-ATPase activity (Kane, 1995; McGuire & Forgac, 2018; Ratto et al., 2022; Stransky & Forgac, 2015). Except for VHA-18 (V1 H subunit), we were able to detect all other subunits of lysosomal v-ATPase, including VHA-5, 6, 7 and UNC-32 (V0 a subunits), VHA-1, 2, 3 and 4 (V0 c subunits), VHA-16 (V0 d subunit), VHA-17 (V0 e subunit), VHA-13 (V1 A subunit), VHA-12 (V1 B subunit), VHA-11 (V1 C subunit), VHA-14 (V1 D subunit), VHA-8 (V1 E subunit), VHA-9 (V1 F subunit), VHA-10 (V1 G subunit), and VHA-15 (V1 H subunit), and also two v-ATPase transporting accessory proteins, VHA-19 and VHA-20 (Figure 2D, Supplementary Table 1). Among the V0 domain subunits, VHA-4, 5, 6, 7 and 16 and UNC-32 are enriched over 10-fold in all four replicates, VHA-1, 2 and 3 are enriched over 10-fold in three replicates and over 5-fold in one replicate, and the low abundant VHA-17 was only detected in two replicates, with more than 10-fold enrichments in both (Figure 2D). The VHA-19 transporting accessory protein is enriched over 10-fold in two replicates and less than 5-fold in two replicates (Figure 2D). In contrast, for the subunits of the V1 domain and the VHA-20 transporting accessory protein, they show no enrichment in the purified lysosomes compared to the flow-through controls (Figure 2D). These results suggest that the free form of the V1 domain and the associated form bound with the V0 domain at lysosomes both exist under the well-fed condition in wild-type worms.

In addition to 30.7% of proteins with known lysosome/endosome localization, the lysosome-enriched proteome includes a small portion of proteins localized to other cellular organelles, ER/Golgi (6.0%), mitochondria (1.4%), peroxisome (0.4%), lipid droplet (0.9%), and synaptic vesicle (1.8%) (Figure 2E). On the other hand, there is a large portion of proteins with annotated plasma membrane localization (24.8%) (Figure 2E). Many of these plasma membrane proteins are receptors that are known to be subject to endocytosis and subsequent recycling lysosomal degradation, such as INA-1/integrin alpha-6 (De Franceschi et al., 2015), VER-3/vascular endothelial growth factor receptor (Ewan et al., 2006), PTC-1/protein patched receptor (Gallet & Therond, 2005), and IGLR-2/leucine-rich repeat-containing G-protein coupled receptor (Snyder et al., 2013) (Supplementary Table 1) (Braulke & Bonifacino, 2009). We also identified proteins involved in the endocytosis process, including low-density lipoprotein receptor-related proteins, LRP-1 (Grant & Hirsh, 1999) and arrestin domain-containing proteins, ARRD-13 and ARRD-18 (Kang et al., 2014) (Supplementary Table 1) that mediate the internalization of plasma membrane receptors (Ma et al., 2002). Thus, the lysosome-enriched proteome also reveals membrane receptor proteins that undergo recycling through the endo-lysosomal system.

### Lysosome-enriched proteome alterations associate with longevity

Lysosomes are known as a heterogeneous population of vesicles, differing in their size, shape, pH and cellular distribution. Considering the emerging role of lysosomes as a cellular hub to integrate protein signals and regulate longevity, we expect that the protein composition of lysosomes may exhibit heterogeneity in association with different longevity mechanisms. To test this hypothesis, we crossed LMP-1 *LysoTg* with the long-lived *lipl-4* transgenic strain (*lipl-4 Tg*) that constitutively expresses a lysosomal acid lipase (Wang et al., 2008) and also the long-lived loss-of-function mutant of *daf-2* (*daf-2(lf)*) that encodes the insulin/IGF-1 receptor (Kenyon et al., 1993; Martins et al., 2016). Reducing insulin/IGF-1 signaling is a well-conserved mechanism to promote longevity in diverse organisms ranging from *C. elegans, Drosophila,* mice to humans (Kenyon et al., 1993; Martins et al., 2016). We conducted Lyso-IP proteomic analyses and compared lysosome-enriched proteomes between wild-type and long-lived strains. The correlation analysis shows good reproducibility among three independent biological replicates (Supplementary Figure 1C, 1D), and the PCA analysis shows a clear separation between Lyso-IP replicates and flow-through controls (Supplementary Figure 1E).

Using LMP-1 Lyso-IP, we have identified 449 lysosome-enriched proteins in the *lipl-4 Tg* worms (Supplementary Table 2), and 176 of them overlap with the LMP-1 Lyso-IP candidates from WT worms (Figure 3A). Thus, 82% of proteins enriched on WT lysosomes are also enriched on *lipl-4 Tg* lysosomes; however, 61% of proteins enriched on *lipl-4 Tg* lysosomes are absent in WT lysosomes (Figure 3A). Interestingly, among the proteins that are specific to *lipl-4 Tg* lysosomes, there are autophagosome proteins and proteins that mediate the fusion between autophagosomes and lysosomes, including ATG-9/ATG9A(Popovic & Dikic, 2014), EPG-7/RB1CC1 (Nishimura et al., 2013), VAMP-7/VAMP8 (Diao et al., 2015; Itakura et al., 2012) and Y75B8A.24/PI4KIIα (Sugiura et al., 1989) (Figure 3B), suggesting that autophagy is induced by *lipl-4 Tg*. Consistently, the induction of autophagy has been reported in the *lipl-4 Tg* worms, which is required for the longevity effect (Lapierre et al., 2011).

**Figure 3.**
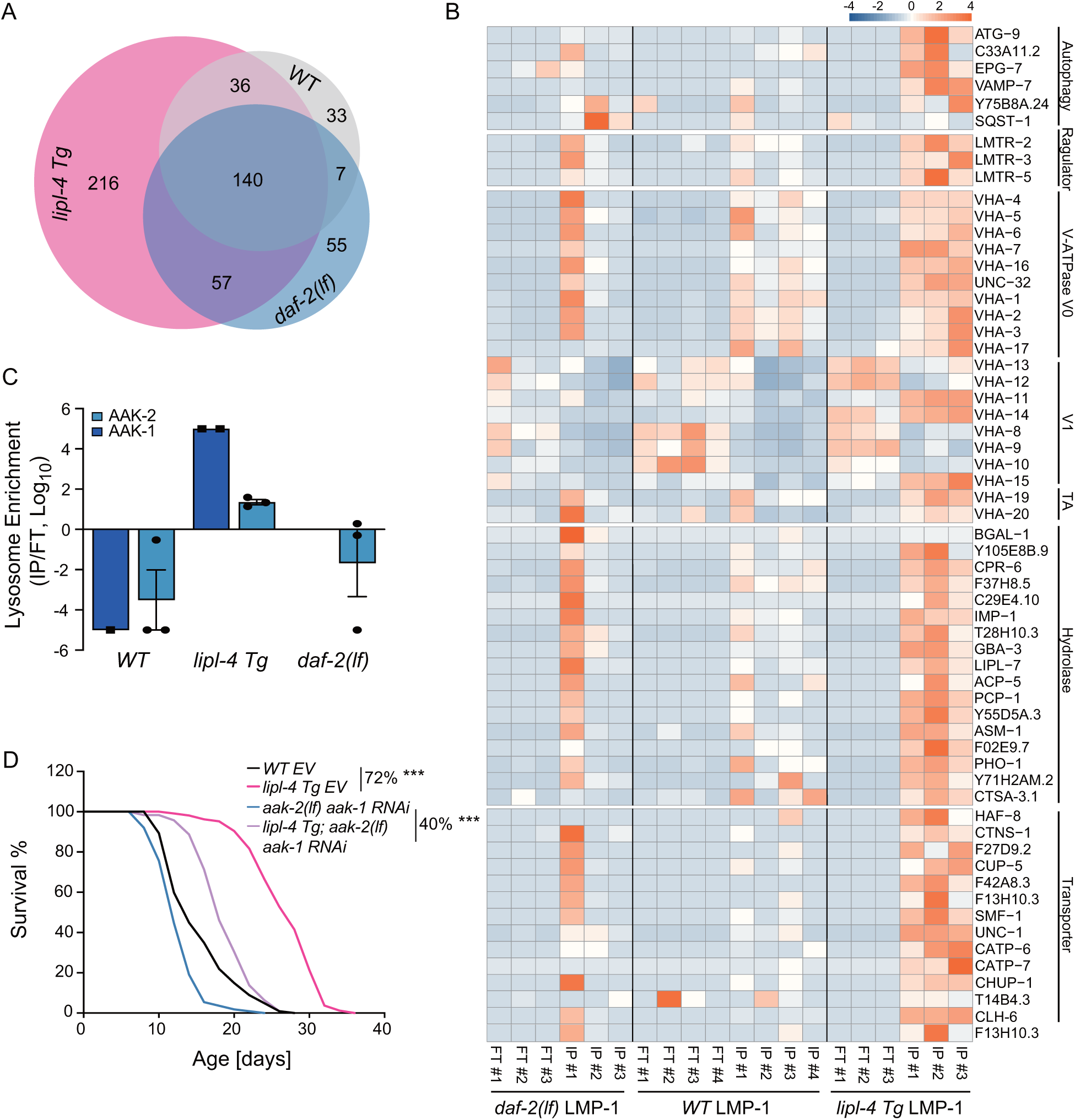
Lysosome-enriched proteome associated with longevity. (**A**) Venn diagram showing overlaps among lysosome-enriched proteomes from wild-type (WT), *lipl-4 Tg* and *daf-2(lf)* worms. (B) Normalized protein levels (z-score across samples) of autophagy-related components, mTORC1 signaling factors, lysosomal v-ATPase V0, V1 and transporting accessory (TA) subunits, lysosomal hydrolases and transporter proteins from LMP-1 Lyso-IP proteomic analyses of WT, *lipl-4 Tg* and *daf-2(lf)*) worms. (C) The lysosomal enrichment ratio (Lyso-IP vs FT) for two homologs of AMPK catalytic subunits, AAK-1 and AAK-2 in WT, *lipl-4 Tg* and *daf-2(lf)*) worms. (D) Reduction of AMPK using the loss-of-function mutant of *aak-2, aak-2(lf)* together with *aak-1* RNAi knockdown reduces the lifespan extension in the *lipl-4 Tg* worms from 72% to 40%. *** *p<0.001* by Log-rank test. The lifespan data are also in supplementary table 3.

In addition, we found that the Ragulator complex, LMTR-2/LAMTOR2, LMTR-3/LAMTOR3 and LMTR-5/LAMTOR5, that serves as a scaffold for the activation of mTORC1 and AMPK (Zhang et al., 2014) is more specifically enriched at the lysosome from the *lipl-4 Tg* worms than WT (Figure 3B). There are two homologs of AMPK catalytic units, AAK-1 and AAK-2 in *C. elegans.* We found that AAK-2 is enriched more than 10-fold in the Lyso-IP samples compared to the flow-through controls from the *lipl-4 Tg* worms, but it is only detected in the flow-through controls from WT worms (Figure 3C). Likely due to its low abundance, AAK-1 was detected twice in the Lyso-IP samples from the *lipl-4 Tg* worms but once only in the flow-through sample from WT worms (Figure 3C). These results suggest that AMPK is specifically recruited to the lysosomal surface and becomes activated to promote longevity in the *lipl-4 Tg* worms. In supporting this idea, we found that the inactivation of AMPK using the *aak-2* loss-of-function mutant together with the *aak-1* RNA interference (RNAi) knockdown reduces the lifespan extension in the *lipl-4 Tg* worms from 72% to 40% (Figure 3D, Supplementary Table 3). Thus, the lysosomal activation of AMPK responds to LIPL-4-induced lysosomal lipolysis and mediates the longevity regulation.

Moreover, compared to WT lysosomes, the enrichment of lysosomal v-ATPase is higher in *lipl-4 Tg* lysosomes, especially for the V0 subunits, VHA-1, 2, 3 the V1 subunits, VHA-11 and VHA-15, and the v-ATPase transporting accessory proteins, VHA-19 and VHA-20 (Figure 3B). There are also 13 lysosomal channels/transporters, including T14B4.3/ATP6AP2, the proton-translocating ATPases required for the v-ATPase-mediated lysosomal acidification (Cruciat et al., 2010) and CLH-6/CLCN7, the H(+)/Cl(-) exchange transporter mediating the acidification of the lysosome (Graves et al., 2008; Nicoli et al., 2019), and 16 lysosomal hydrolases that are specifically associated with the *lipl-4 Tg* lysosomes (Figure 3B). Together, these results suggest that the proportion of mature lysosomes is increased in the *lipl-4 Tg* worms, which may lead to increased autophagy, the lysosomal activation of AMPK, and consequently the induction of longevity.

In parallel, 259 lysosome-enriched proteins were identified in the *daf-2(lf)* mutant using LMP-1 Lyso-IP (Supplementary Table 4), 147 of them overlapping with the LMP-1 Lyso-IP candidates from WT worms, 197 of them overlapping with the LMP-1 Lyso-IP candidates from the *lipl-4 Tg* worms, and 55 unique to the *daf-2(lf)* mutant (Figure 3A). The *daf-2(lf)* lysosome-enriched proteome shows enrichment of autophagic components, including ATG-9/ATG9A (also shown in *lipl-4 Tg* lysosome-enriched proteome) and the autophagy receptor SQST-1/SQSTM1 (Figure 3B), which supports the previously reported induction of autophagy in the *daf-2(lf)* mutant. On the other hand, the Ragulator complex components LMTR-2/3/5 were not more enriched in the lysosome from the *daf-2(lf)* mutant than that from WT worms (Figure 3B). For the AMPK catalytic subunits, AAK-2 was not identified in the lysosome-enriched proteome from the *daf-2(lf)* mutant, and the low abundant AAK-1 was not detected in either Lyso-IP or flow-through samples (Figure 3C). It is known that the activation of AMPK displays high spatial specificity in the cell when responding to different upstream stimuli (Khan & Frigo, 2017). In *C. elegans,* it was previously shown that AAK-2 mediates the longevity effect conferred by the *daf-2(lf)* mutant (Apfeld et al., 2004). Our results indicate that this regulation might not be associated with the lysosomal activation of AMPK. Thus, the spatial specificity of AMPK activation at different subcellular compartments may be linked with different longevity mechanisms.

### Enhanced lysosome-nucleus interaction mediates longevity

In our previous studies, we found that upon LIPL-4-induced lysosomal lipolysis, a lysosome-to-nucleus retrograde lipid messenger signaling pathway is activated in intestinal cells to promote longevity (Folick et al., 2015; Ramachandran et al., 2019; Savini et al., 2022; Wang et al., 2008). Interestingly, when analyzing the LMP-1 lysosome-enriched proteome in the *lipl-4 Tg* worms, we found an enrichment of nucleus-localized proteins (Figure 4A), including two nucleoporin proteins NPP-6/Nup160 and NPP-15/Nup133 in the Nup160 complex that localizes at the basket side of the nuclear pore (Figure 4B) (Vasu et al., 2001). Such enrichment of nucleoporin proteins was not found in the LMP-1 lysosome-enriched proteome of either WT or the *daf-2(lf)* mutant (Figure 4B). In the cell, mobile lysosomes change their distribution along the perinuclear-peripheral axis in response to different nutrient signals and metabolic status (Ballabio & Bonifacino, 2020; Pu et al., 2016). Moreover, it is known that the cellular position of lysosomes affects their luminal pH, with perinuclear lysosomes being more acidic (Johnson et al., 2016). We found that the lysosome-enriched proteome of the *lipl-4 Tg* worms consists of lysosomal v-ATPase V1 subunits, proton-translocating ATPase and exchanging ion transporters that mediate lysosomal acidification, but these components were not enriched in the WT lysosome proteome (Figure 3B), supporting that lysosomes are more acidic in the *lipl-4 Tg* worms. We thus hypothesize that LIPL-4-induced lysosomal lipolysis may enhance the interaction between lysosomes and the nucleus and further facilitate the lysosome-to-nucleus retrograde lipid signaling to promote longevity.

**Figure 4.**
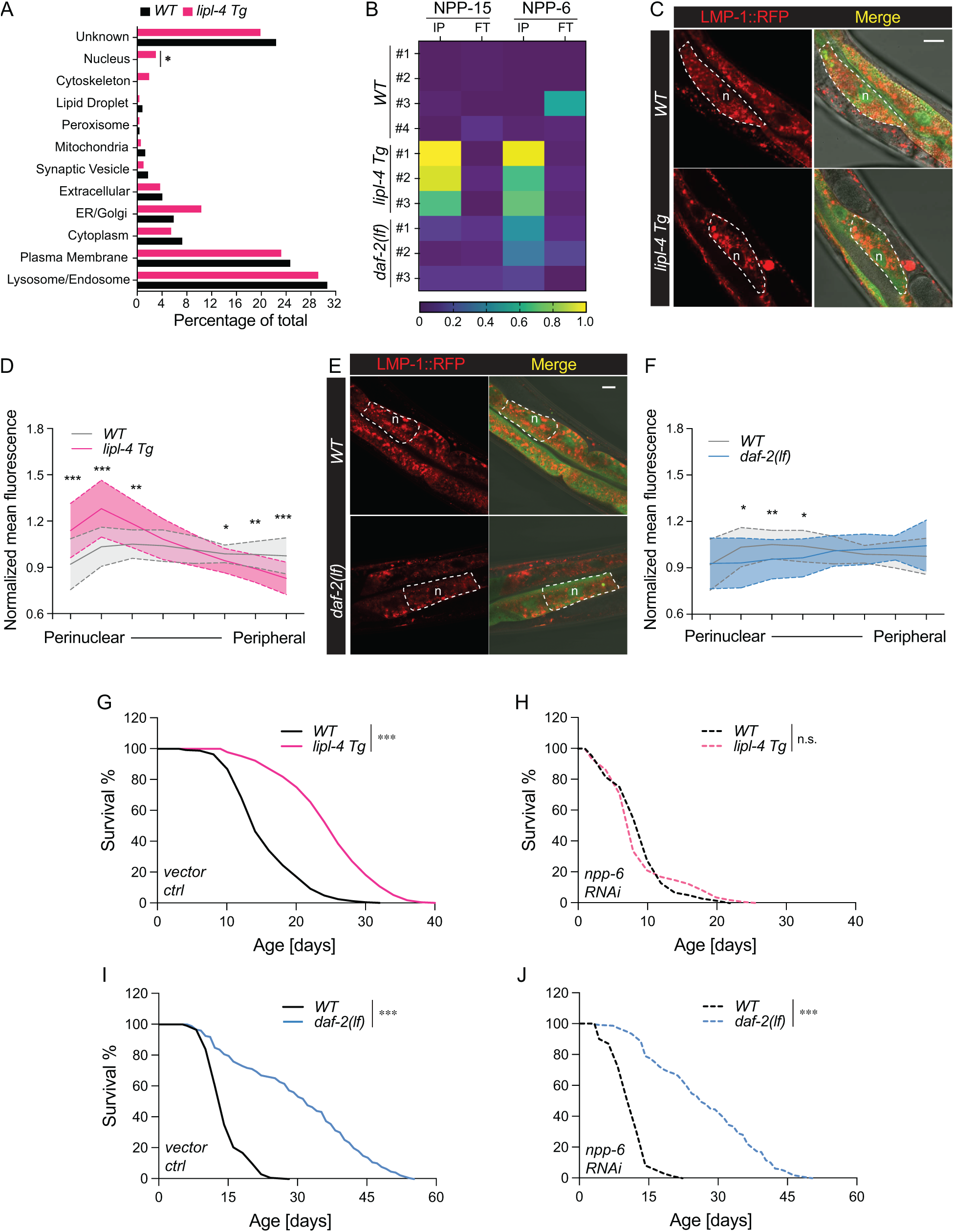
Enhanced lysosome-nucleus interaction mediating longevity. (**A**) The percentage of proteins with different subcellular localization is compared between lysosome-enriched proteomes from WT and *lipl-4 Tg* worms. *\* p=*0.019 by 2-sample test for equality of proportions. **(B)** Heatmap showing the average levels of nucleoporin proteins NPP-6 and NPP-15 in Lyso-IP (IP) and flow-through (FT) samples from WT, *lipl-4 Tg* and *daf-2(lf)* worms. **(C, E)** Representative images of intestinal cells in WT, *lipl-4 Tg* (**C**) and *daf-2(lf)* (**E**) worms carrying LMP-1::RFP-3×HA and nucleus-enriched GFP, showing the accumulation of lysosomes around the perinuclear region in the *lipl-4 Tg* but not *daf-2(lf)* worms. Dashed lines circle intestinal cells and n marks the nucleus. Scale bar=20 μm. **(D, F)** Line graph showing the spatial distribution of lysosomes from the nuclear to peripheral region quantified by normalized regional RFP fluorescence signals in intestinal cells of WT, *lipl-4 Tg* (**D**) and *daf-2(lf)* (**F**) worms. N =50 WT /33 *lipl-4 Tg*, 33 WT/ 28 *daf-2(lf*). Data are represented as mean ± SD. *p* values for (D) (from left to right): 1.23x10^-7^, 2.25x10^-5^, 0.00322, 0.368, 0.273, 0.0447, 0.00268, 1.20x10^-5^; *p* values for (F) (from left to right): 0.633, 0.0211, 0.00259, 0.0359, 0.767, 0.151, 0.106, 0.0671. **(G-H)** *lipl-4 Tg* worms show lifespan extension compared to WT worms (**G**), which is fully suppressed by RNAi knockdown of *npp-6* (**H**). \*\*\**p<0.001,* n.s. *p>0.05* by Log-rank test. **(I-J)** *daf-2(lf)* worms show lifespan extension compared to WT worms (**I**), which is not affected by *npp-6* RNAi knockdown (**J**). \*\*\**p<0.001* by Log-rank test. The lifespan data are also in supplementary table 3.

To test this hypothesis, we imaged lysosomal positions in intestinal cells where *lipl-4* is expressed. Using a dual reporter strain expressing both lysosomal LMP-1::RFP fusion and nucleus-localized GFP, we found that lysosomes exhibit a dispersed pattern in the intestinal cell of WT worms (Figure 4D). However, in the *lipl-4 Tg* worms, lysosomes are clustered in the perinuclear region (Figure 4D), supporting the hypothesis that the lysosome-nucleus interaction is enhanced. To quantitatively measure this change in lysosomal positioning, we analyzed the RFP fluorescent signal distribution in intestinal cells (Supplementary Figure 2A). We found the perinuclear and peripheral distribution of lysosomes in the *lipl-4 Tg* worms is significantly increased and decreased, respectively, compared to WT worms (*p<0.01*, Figure 4C, 4D, Supplementary Figure 2B). In contrast, such perinuclear clustering is not observed in intestinal cells of the *daf-2(lf)* mutant (Figure 4E, 4F, Supplementary Figure 2C). Moreover, we found that the RNAi knockdown of *npp-6* suppresses the lifespan extension in the *lipl-4 Tg* worms (Figure 4G, 4H) but does not affect the lifespan extension in the *daf-2(lf)* mutant (Figure 4I, 4J). These results suggest that the enhanced lysosome-nucleus interaction is specifically related to LIPL-4-induced lysosomal signaling and longevity.

Together, these studies discover changes in the lysosome-enriched proteome between WT and different long-lived strains and reveal that the lysosomal proteome composition can be influenced by the metabolic status of the lysosome and its interaction with other organelles and in turn correlates with lysosomal signaling and longevity regulation.

### Profile lysosome-enriched proteome heterogeneity among different tissues

Tissue-specific regulation of longevity has been reported in various organisms. To examine how lysosome-enriched proteomes exhibit heterogeneity among different tissues, we have generated four transgenic strains that express LMP-1 Lyso-Tag specifically in neurons, muscle, intestine and hypodermis using tissue-specific promoters, *unc-119*, *myo-3*, *ges-1* and *col-12*, respectively (Figure 5A). Using these transgenic strains, we purified lysosomes in a tissue-specific manner and conducted proteomic profiling. The correlation analysis shows good reproducibility among three independent biological replicates (Supplementary Figure 3A-D).

**Figure 5.**
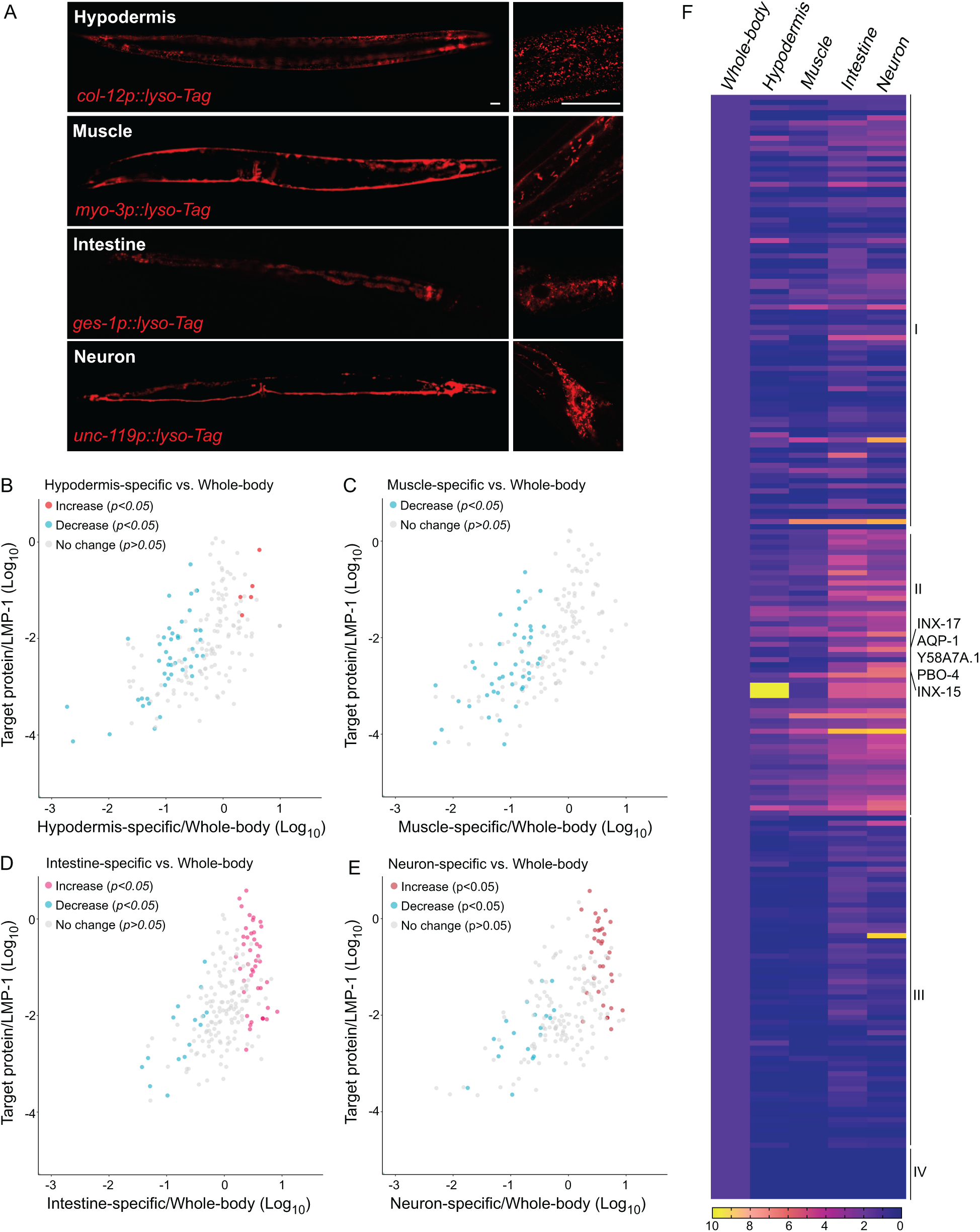
Lysosomal proteome heterogeneity across tissues. (**A**) Example images of transgenic strains carrying Lyso-Tag (LMP-1::RFP-3×HA) driven by tissue-specific promoters. Scale bar=20 μm. **(B-E)** Scatter plot showing the relative enrichment ratio for each of 216 lysosome-enriched proteins identified from whole-body LMP-1 Lyso-IP in comparison with tissue-specific LMP-1 Lyso-IPs, hypodermis (**B**), muscle (**C**), intestine (**D**) and neuron (**E**). X axis, enrichment ratio tissue-specific vs. whole-body; Y axis, normalized protein abundance over LMP-1; each dot represents the average of three replicates. **(F)** Heatmap showing the relative enrichment of 216 lysosome-enriched proteins identified from whole-body LMP-1 Lyso-IP in comparison with tissue-specific LMP-1 Lyso-IPs. Group I, comparable ratios between whole-body and tissue-specific Lyso-IPs; Group II, increase in tissue-specific Lyso-IPs (*p<0.05* by student’s t-test); Group III, decrease in tissue-specific Lyso-IPs (*p<0.05* by student’s t-test); Group IV, absent in tissue-specific IPs.

Unlike the whole-body Lyso-IP, the flow-through samples from tissue-specific Lyso-IP contain not only non-lysosomal fractions from the targeted tissue but also lysosomes from non-targeted tissues. Thus, these flow-through samples cannot be simply used as controls to determine the enrichment of proteins at the lysosome in the targeted tissue. To assess tissue-specific changes, we have normalized the level of each identified protein to the level of LMP-1 in the same replicate, and then compared the normalized ratio between the whole-body Lyso-IP and the tissue-specific Lyso-IP (Figure 5B-E, Supplementary Table S5). We found that among the 216 proteins identified from the whole-body Lyso-IP, 85 of them show comparable ratios between the whole-body Lyso-IP and the four tissue-specific Lyso-IPs (Figure 5F, Group I), suggesting relative homogenous lysosomal enrichments of these proteins among different tissues. Nine of them were completely absent in the tissue-specific Lyso-IP, which may be related to their low abundance (Figure 5F, Group IV). There are also proteins that were not detected in one or two tissues but exhibited similar enrichments as the whole body Lyso-IP in other tissues, such as PGP-6, an ABC transporter missing in the muscle-specific Lyso-IP, PBO-1, calcium-binding protein P22 missing in the hypodermis-specific Lyso-IP and Y57G11C.33, Golgin A7 missing in both the hypodermis- and neuron-specific Lyso-IPs, which may be associated with their lack of expression and/or lack of lysosome enrichments in specific tissues.

Importantly, there are 122 proteins that exhibited significant difference in their enrichments between the whole-body Lyso-IP and the tissue-specific Lyso-IPs (*p<0.05*), 56 of them (Group II) showing an increase in the tissue-specific Lyso-IPs while the other 66 (Group III) showing a decrease (Figure 5F). Some of them were only enriched in one tissue but remained unchanged in others, including 15 proteins with higher ratios only in the intestine, 3 only in the hypodermis and 5 only in neurons. There are also five proteins that showed an increased enrichment in one tissue, but a decreased enrichment in another tissue, including AQP-1, an Aquaporin channel with a higher ratio in neurons but a lower one in the hypodermis, Y58A7A.1 a copper uptake transporter with a higher ratio in the hypodermis but a lower one in the muscle, PBO-4, a sodium/hydrogen exchanger with a higher ratio in the intestine but a lower one in the hypodermis, and two innexin proteins that form gap junctions in invertebrate animals, INX-17 with a higher ratio in neurons but a lower ratio in the hypodermis and INX-15 with a higher ratio in the intestine but lower ratios in the muscle and hypodermis (Figure 5F). For these candidates, there is a clear tissue preference regarding their lysosomal enrichments. Thus, the lysosomal proteome is not only affected by the genotype of the organism, but also exhibits heterogeneity among different tissues within the organism, which may be related to the metabolic status in each tissue and consequently contribute to specific activities and signaling effects of the tissue. Our studies provided a list of candidates for further investigation of the tissue-specific regulation of lysosomal metabolism and signaling.

### Cystinosin positive mature lysosomes enrich specific lysosomal proteins

The analysis of the candidates specifically detected in the *lipl-4 Tg* worms suggest that the proportion of mature lysosomes may affect the lysosomal protein composition. Although LMP-1 is a well-established lysosomal protein marker and highly abundant on the lysosomal surface, it can be also detected in late endosomes and sometimes in early endocytic compartments. With the hope to profile proteins enriched in mature lysosomes, we chose CTNS-1, the *C. elegans* lysosomal cystine transporter Cystinosin that is a well-established marker of mature lysosomes (Gahl et al., 1982; Jonas et al., 1982; Kalatzis et al., 2001), and generated a transgenic strain expressing CTNS-1 tagged with both 3×HA and RFP (CTNS-1 *lysoTg*). Fluorescence imaging of RFP confirmed the lysosomal localization of the CTNS-1 fusion protein in live organisms (Figure 6A). Using this transgenic strain, we followed the same Lyso-IP and MS profiling pipeline. The correlation analysis shows good reproducibility among three independent biological replicates (Supplementary Figure 1G), and the PCA analysis indicates a clear separation between Lyso-IP samples and flow-through controls (Supplementary Figure 1F).

**Figure 6.**
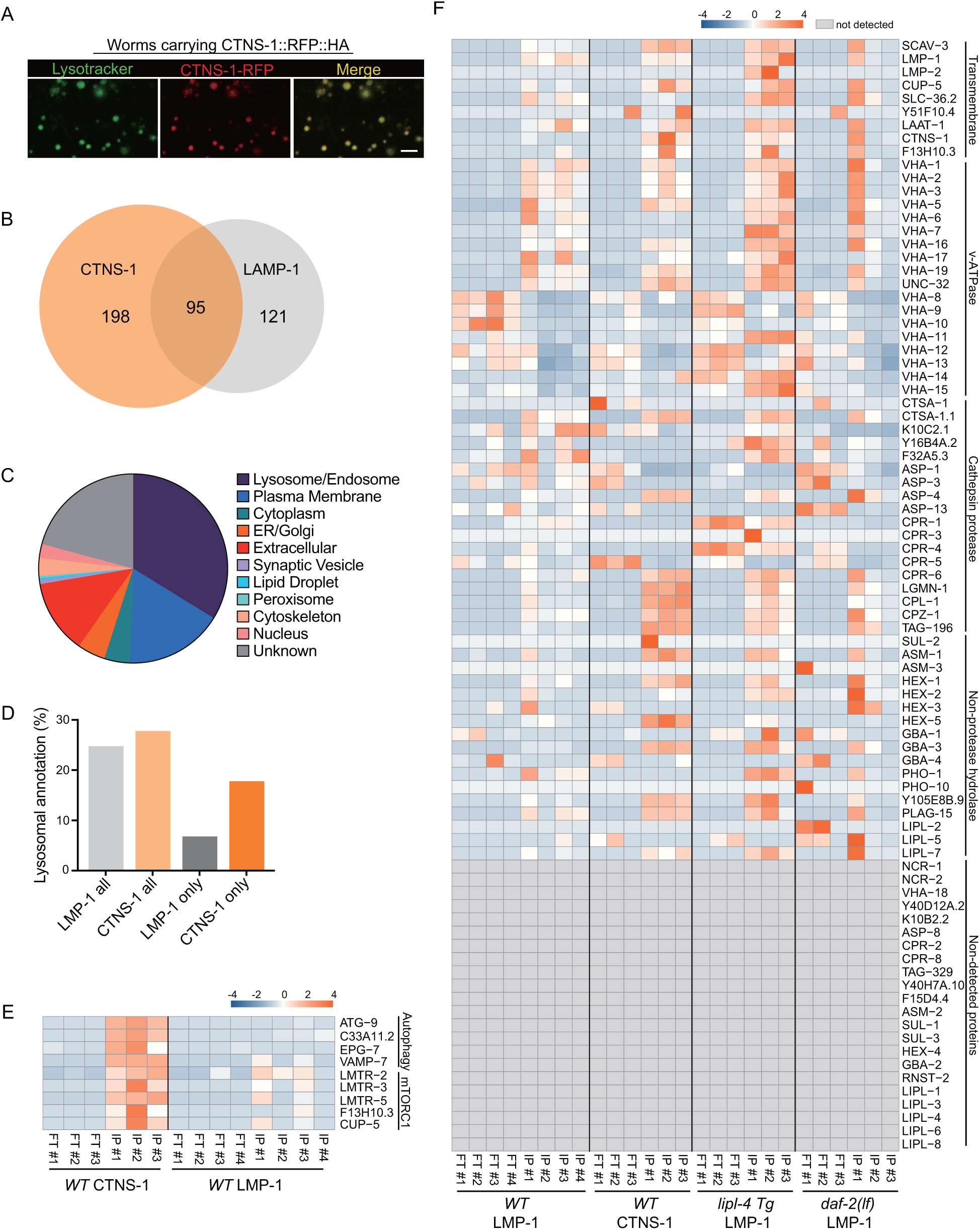
Lysosome-enriched proteome identified with Cystinosin. (**A**) Example images of transgenic strains carrying CTNS-1 Lyso-Tag (CTNS-1::RFP-3×HA) with LysoTracker staining to mark lysosomes *in vivo*. Scale bar=5 µm. **(B)** Venn diagram showing the overlap between lysosome-enriched proteomes using LMP-1 Lyso-IP and CTNS-1 Lyso-IP. **(C)** Pie chart showing subcellular location categories of lysosome-enriched proteins. **(D)** The proportion of candidates with lysosomal localization annotation in different candidate groups. “LMP-1 all” and “CTNS-1 all”, all candidates from LMP-1 Lyso-IP and CTNS-1 Lyso-IP, respectively; “LMP-1 only” and “CTNS-1 only”, candidates only identified from LMP-1 Lyso-IP or CTNS-1 Lyso-IP, respectively. **(E)** Normalized protein levels (z-score across samples) of autophagy-related components and mTORC1 signaling factors from LMP-1 and CTNS-1 Lyso-IP proteomic analyses of WT worms. **(F)** Normalized protein levels (z-score across samples) of previously annotated lysosomal proteins from LMP-1 Lyso-IP proteomic analyses of WT, *lipl-4 Tg* and *daf-2(lf)*) worms and CTNS-1 Lyso-IP proteomic analyses of WT worms. Proteins not detected by proteomic profiling are marked in grey.

Using the same selection criteria, we identified 293 candidates whose levels are enriched at least 10-fold in the purified lysosomes than those in the flow-through controls among all independent biological replicates and show over 2-fold enrichment compared to the non-tag controls (Supplementary Table 6). There are 95 lysosome-enriched proteins shared between the LMP-1 and the CTNS-1 Lyso-IP proteomic profiling datasets (Figure 6B, Supplementary Table 6), and 47 of these shared proteins are annotated with lysosomal localization (Supplementary Table 6). We have also crossed the CTNS-1 *lysoTg* strain with the *lipl-4 Tg* strain and the *daf-2(lf)* mutant and then conducted Lyso-IP proteomic profiling. However, the pull-down efficiency was very low in these long-lived worms, which prevents us to identify proteins unique to CTNS-1 Lyso-IP in those conditions.

In WT worms, the proportions of the identified proteins with different categories of subcellular annotation are comparable between LMP-1 and CTNS-1 Lyso-IP conditions (Figure 6C, 2E), and for proteins with lysosomal annotation, the proportion is 25% and 28% in LMP-1 and CTNS-1 Lyso-IP, respectively (Figure 6D). However, among the 121 proteins only identified in LMP-1 Lyso-IP, there are only 8 with lysosomal annotation (7%); while for the 198 proteins only identified in CTNS-1 Lyso-IP, 35 are with lysosomal annotation and the proportion remains as 18% (Figure 6D). Among the lysosomal proteins that are unique to CTNS-1 Lyso-IP, there are autophagosome proteins and proteins that mediate the fusion between autophagosomes and lysosomes, including ATG-9/ATG9A (Popovic & Dikic, 2014), C33A11.2/DRAM2 (Crighton et al., 2006), EPG-7/RB1CC1 (Nishimura et al., 2013), and VAMP-7/VAMP8 (Diao et al., 2015; Itakura et al., 2012) (Figure 6E). CTNS-1 Lyso-IP unique candidates also include the Ragulator complex components LMTR-2/3/5, the lysosomal amino acid transporter F13H10.3/SLC38A9 and the lysosomal calcium channel CUP-5/TRPML1 that regulate mTORC1 signaling (Li et al., 2016; Rebsamen et al., 2015; Wang et al., 2015; Wyant et al., 2017) (Figure 6E). These results suggest that CTNS-1 Lyso-IP enriches lysosomes that are actively involved in autophagy and transduce mTORC1 signaling.

Furthermore, when systemically examining 85 lysosome-related proteins that were previously annotated in *C. elegans* based on sequence homology (Sun et al., 2020), we found that many lysosomal hydrolases exhibit increased enrichments with CTNS-1 Lyso-IP, especially for those cysteine proteases including CPR-6, LGMN-1, CPL-1, CPZ-1 and TAG-196 (Figure 6F). These enrichments are consistent with that CTNS-1 is located at mature lysosomes as a cysteine transporter (Gahl et al., 1982; Jonas et al., 1982; Kalatzis et al., 2001). The similar increased enrichments of lysosomal hydrolases were also observed in the LMP-1 Lyso-IP candidates identified from the *lipl-4 Tg* worms (Figure 6F), further supporting that the long-lived *lipl-4 Tg* worms carry more acidic lysosomes. Together, we found that lysosome-enriched proteomes identified from both LMP-1 and CTNS-1 Lyso-IP consist of well-characterized lysosomal enzymes and integral membrane proteins as well as proteins that contribute to lysosomal signaling, dynamics and contact with other cellular compartments. Besides many known lysosomal proteins, various proteins that are not previously linked with lysosomes are now identified through these systemic analyses.

### Lysosome-enriched proteins regulate different lysosomal activities

To understand the role of these newly identified lysosome-enriched proteins in regulating lysosomal functions, we have examined their effects on lysosomes using an RNAi screen based on LysoSensor fluorescence intensity. We focused on 95 lysosome-enriched proteins shared between LMP-1 and CTNS-1 Lyso-IPs and knocked down their coding genes by RNAi, and then used LysoSensor probes to stain lysosomes. From screening these 95 candidates (Supplementary Table 7), we have identified five genes whose inactivation cause changes in LysoSensor signal intensity, and four of them have human homologs, including two lysosomal v-ATPase subunits, UNC-32/ATP6V0A and VHA-5/ATP6V0A, the lysosomal amino acid transporter SLC-36.2/SLC36A1 (SLC36A4), and a transmembrane protein R144.6/TMEM144 (Supplementary Table 7, Supplementary Figure 5). We further examined their effects on the lysosomal number, size and pH. We found that the RNAi knockdown of the two lysosomal v-ATPase subunits, UNC-32 and VHA-5, lead to decreased lysosomal numbers (Figure 7F), but an increase in the lysosomal size (Figure 7G). Using fluorescence lifetime microscopy, we also found that the RNAi knockdown of R144.6 and UNC-32 increases and decreases lysosomal pH, respectively (Figure 7H).

**Figure 7.**
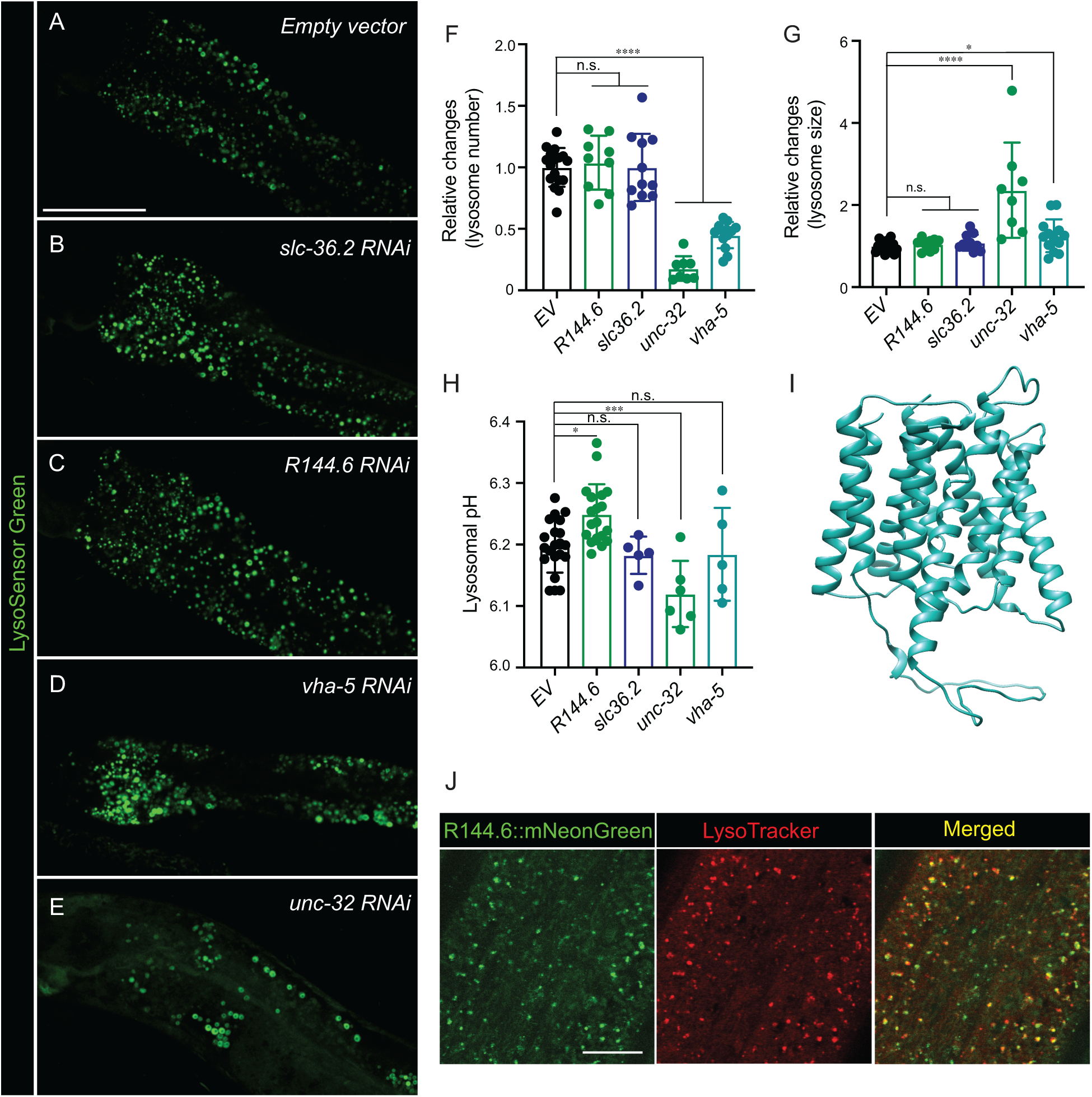
Lysosome-enriched proteins regulating lysosomal functions. (**A-E)** Confocal fluorescence microscopy images of intestinal cells in worms stained with LysoSensor DND-189 and treated with *empty vector* (**A**), *slc36.2* RNAi (**B**), *R144.6* RNAi (**C**), *vha-5* RNAi (**D**) and *unc-32* RNAi (**E**). Scale bar=50 μm. **(F, G)** RNAi knockdown of *unc-32* or *vha-5* decreases the lysosome number *(****p< 0.0001)* (**F**) but increases the lysosome size *(****p< 0.0001, *p=0.0222)* (**G**). The average lysosome number and size per pair of intestinal cells were quantified. Data are shown as mean ± SD. Student’s t-test (unpaired, two-tailed) was performed between the *empty vector* group and each RNAi-treated group. n = 17(*EV*), 9(*R144.6*), 11(*slc36.2*), 8(*unc-32*), 13(*vha-5*). n.s. *p>0.05,* **(H)** RNAi knockdown of *R144.6 (*p= 0.0206)* and *unc-32 (***p= 0.0001)* increase and decrease lysosomal pH, respectively. Lysosomal pH was calculated based on LysoSensor’s lifetime measured by Fluorescence Lifetime Microscopy. Data are shown as mean ± SD. Student’s t-test (unpaired, two-tailed) was performed between the *empty vector* group and each RNAi-treated group. n = 21(*EV*), 19(*R144.6*), 5(*slc36.2*), 6(*unc-32*), 5(*vha-5*). n.s. *p>0.05*. **(I)** The structure of the R144.6 protein predicted by AlphaFold2 supports it as a solute carrier family transporter. **(J)** Confocal fluorescence microscopy images show that mNeonGreen signals from endogenously tagged R144.6 colocalize with LysoTracker Red signals in the hypodermis. Scale bar=10 μm.

UNC-32, VHA-5 and SLC-36.2 are well-known lysosomal proteins, but the subcellular localization of the R144.6 protein remains unknown. R144.6 is a predicted carbohydrate transporter, and structural simulation using AlphaFold2 suggested it as a solute carrier family (SLC) transporter (Figure 7I). To characterize its subcellular localization, we have generated a CRISPR knock-in line in which the endogenous R144.6 protein is fused with the fluorescent protein NeonGreen and then stained these worms with LysoTracker Red to mark lysosomes. In the hypodermis, we found that NeonGreen and LysoTracker Red signals overlap, confirming the lysosomal localization of this newly identified transmembrane protein from Lyso-IP (Figure 7J). On the other hand, its expression was not detected in the muscle, and in the intestine, its NeonGreen signals were not overlapped with LysoTracker Red, which is consistent with tissue-specific Lyso-IP analyses. R144.6 were enriched in the hypodermis-specific Lyso-IP at a similar level as in the whole-body Lyso-IP; however, it was not detected in the muscle- or intestine-specific Lyso-IP (Supplementary Table 5). These results further support the enrichment specificity of proteins at the lysosome in different tissues as well as the power of the Lyso-IP proteomic profiling in discovering new lysosomal proteins with functional significance.

## DISCUSSION

Our studies reveal the heterogeneity of lysosomal protein composition that is associated with lysosomal status, tissue specificity, and organism longevity. Through systemic profiling of lysosome-enriched proteins under different conditions, we confirmed the induction of lysosome-related autophagy by different longevity-promoting pathways, unveiled increased cellular interaction between lysosomes and the nucleus upon the induction of lysosomal lipolysis and its contribution to longevity regulation, and underlined the importance of the spatial control of AMPK activation in regulating longevity. Our studies provide not only methods for future studies to profile the dynamics of the lysosomal proteome in response to diverse physiological inputs, but also resources for understanding the vital contribution of these dynamics in modulating signal transduction, organelle crosstalk and organism longevity.

These proteomic studies reveal changes in the interaction between lysosomes and other organelles under different conditions. One example is the lysosome-nucleus interaction. In the Lyso-IP fraction from WT worms, we did not detect any proteins with sole nuclear localization; however, in the Lyso-IP fraction from the *lipl-4 Tg* or *daf-2(lf)* mutant worms, nuclear proteins were identified and the percentage of the increase over WT is significantly higher in the *lipl-4 Tg* worms (*p<0.05*, Figure 4A, Supplementary Figure 4). Based on this finding, we discovered the previously unknown perinuclear accumulation of lysosomes in the *lipl-4 Tg* worms (Figure 4C, 4D) and further confirmed its importance for longevity regulation (Figure 4G, 4H). It is shown previously that perinuclear lysosomes are more acidic than peripheral lysosomes (Johnson et al., 2016; Webb et al., 2021). Thus, the increase in perinuclear lysosomes may be associated with the increased proportion of mature lysosomes in the *lipl-4 Tg* worms, which is supported by the increased enrichments of lysosomal v-ATPase, channels/transporters and hydrolases (Figure 3B). This increased distribution of lysosomes toward the perinuclear region could facilitate proteins and metabolites transporting from the lysosome to the nucleus through the nuclear pore and in turn their signaling effects. On the other hand, we did not detect perinuclear accumulation of lysosomes in the *daf-2(lf)* mutant worms by cellular imaging, and the nuclear proteins detected through LMP-1 Lyso-IP from the *daf-2(lf)* mutant worms are mainly involved in RNA splicing. In yeast cells, defects in pre-RNA processing have been associated with nucleophagy (Leger-Silvestre et al., 2005) that involves SQSTM1 and lysosomes (Ivanov et al., 2013; Mijaljica & Devenish, 2013). We thus speculate that the increased enrichment of nuclear proteins in the *daf-2(lf)* mutant worms may be associated with the induction of nucleophagy but not changes in lysosomal positioning.

mTORC1 and AMPK are key metabolic checkpoints that regulate anabolic and catabolic processes in mutually opposing ways. In sensing the lack of nutrients, AMPK signals activate the catabolic process while inhibiting the anabolic one. On the other hand, responding to nutrient availability, mTORC1 activation upregulates anabolic metabolism and promotes cell growth. Intriguingly, it is now known that both mTORC1 and AMPK are recruited to the lysosomal surface for activation, which requires the scaffold Ragulator complex that consists of LAMTOR subunits. We found that the Ragulator complex (LMTR-2, 3 and 5) is enriched at the lysosome purified from WT worms using CTNS-1 but not LMP-1 Lyso-IP (Figure 6E), suggesting a predominant association of the Ragulator complex with mature lysosomes, which could in turn determine the preference of mTORC1 and AMPK activation at the lysosomal surface. Alternatively, it would be also possible that the Ragulator complex carries a preference toward CTNS-1/Cystinosin-containing lysosomes, which would infer the interaction between lysosomal cysteine metabolism and mTORC1 signaling. Interestingly, previous studies show that Cystinosin co-immunoprecipitates with the Ragulator complex in mammalian cells (Andrzejewska et al., 2016), and in *Drosophila,* cysteine efflux from the lysosome via Cystinosin antagonizes mTORC1 signaling and upregulates the tricarboxylic acid cycle (Jouandin et al., 2022). Whether this inhibitory effect of lysosomal cysteine on mTORC1 is related to the preferential interaction between the Ragulator complex and Cystinosin would be an interesting question for future studies.

Both mTORC1 and AMPK have been implicated in the regulation of longevity across different species, being intertwined with other longevity regulatory mechanisms (Savini et al., 2019). In the long-lived *lipl-4 Tg* worms, the lysosomal enrichment of the Ragulator complex is increased with LMP-1 Lyso-IP, which may be a result of the increased proportion of mature lysosomes upon the induction of lysosomal lipolysis. At the same time, we could not rule out the possibility that the increased enrichment of the Ragulator complex is a result of the induced level of lysosomal CTNS-1/Cystinosin in the *lipl-4 Tg* worms. We found that with LMP-1 Lyso-IP, the level of the CTNS-1/Cystinosin transporter is increased in the *lipl-4 Tg* worms, together with the increase of several cysteine cathepsins (Figure 6F). Our previous studies found that mitochondrial β-oxidation is increased in the *lipl-4 Tg* worms, leading to decreased triglyceride storage (Ramachandran et al., 2019). The *lipl-4 Tg* worms also show an induced autophagy (Mak et al., 2020). These phenotypes are the same as those observed in fruit flies with Cystinosin overexpression (Jouandin et al., 2022). Considering the inhibitory effect of Cystinosin on mTORC1 in fruit flies, the induction of Cystinosin in the *lipl-4 Tg* worms might reduce mTORC1 signaling. In supporting this idea, our unpublished study shows that *lipl-4 Tg* does not further enhance the lifespan extension of the *raga-1* mutant that has reduced mTORC1 signaling. Furthermore, our study reveals the lysosomal enrichment of AMPK in the *lipl-4 Tg* worms, and its requirement for the longevity effect (Figure 3C, 3D). On the other hand, the involvement of lysosomal mTORC1 and AMPK signaling in regulating the longevity effect associated with reduced insulin/IGF-1 signaling was not identified in the *daf-2(lf)* mutant. It would be interesting for future studies to systemically profile lysosome-enriched proteomes in other longevity models, such as caloric restriction, mild inhibition of mitochondrial respiration and germline deficiency, using the same method, which would help understand the involvement of lysosomal metabolism and signaling in these different longevity mechanisms.

## MATERIALS AND METHODS

### *C. elegans* strains and maintenance

The following strains were used in this study: N2, CB1370 *daf-2(e1370),* RB754 *aak-2(ok524)*, *unc-76(e911)*, MCW953 *nre-1(hd20);lin-15b(hd126)*, MCW14 *raxIs3 [ges-1p::lipl-4::SL2GFP]*, MCW859 *raxIs103[sur-5p:lmp-1::RFP-3×HA;unc-76(+)],* MCW935 *daf-2(e1370);raxIs103[sur-5p:lmp-1::RFP-3×HA;unc-76(+)],* MCW923 *raxIs3[ges-1p::lipl-4::SL2GFP];raxIs103[sur-5p:lmp-1::RFP-3×HA;unc-76(+)], MCW861 unc-76(e911);raxEx311[Pmyo-3:lmp-1::RFP-3×HA;unc-76(+)],* MCW924 *unc-76(e911); raxEx346[Pcol-12:lmp-1::RFP-3×HA;unc-76(+)],* MCW862 *unc-76(e911);raxEx312[Punc-119:lmp-1::RFP-3×HA;unc-76(+)],* MCW914 *unc-76(e911);raxEx341[Pges-1:lmp-1::RFP-3×HA;unc-76(+)],* MCW934 *raxIs118[sur-5p:ctns-1::RFP-3×HA;unc-76(+)].* The strains N2, CB1370, and RB754 were obtained from *Caenorhabditis* Genetics Center (CGC). The strain *unc-76(e911)* was obtained from Dr. Zheng Zhou’s Lab. Other strains were generated in our lab.

*C. elegans* strains were maintained at 20°C on standard NGM agar plates seeded with OP50 *E.coli* (HT115 *E. coli* for RNAi experiments) using standard protocols (Stiernagle, 2006) and kept at least three generations without starvation before experiments.

### Molecular cloning and generating transgenics

All the expression constructs were generated using Multisite Gateway System (Invitrogen) as previously described (Mutlu et al., 2020). The *lmp-1* and *ctns-1* coding sequences were PCR-amplified from *C. elegans* cDNA then inframe fused with RFP-3×HA, and all promoters were PCR-amplified from *C. elegans* genomic DNA.

Transgenic strains were generated by microinjecting the day-1-adult germline of *unc-76(e911)* worms with DNA mixture containing expression construct and *unc-76(+)* rescuing plasmid. For integration strains, the stable extrachromosomal arrays were integrated with gamma irradiation (4500 rads for in 5.9 minutes) and backcrossing to wild-type N2 at least 8 times.

### Lysosome immunoprecipitation (Lyso-IP)

Lyso-IP is based on the method used in mammalian cells (Abu-Remaileh et al., 2017). Briefly, transgenic strains stably expressing C-terminal RFP- and 3×HA-tagged lysosomal membrane protein LMP-1 or CTNS-1 under whole-body *Psur-5* or tissue-specific promoters were generated. Around 160,000 day-1-adult worms per genotype were collected, washed 3 times with M9 buffer then washed 1 time with KPBS buffer (136 mM KCl, 10 mM KH_2_PO_4_). Worms in 2 ml KPBS were quickly homogenized with Dounce homogenizer on ice until no visible animals were seen under the microscope. The lysate was centrifuged at 1000 g for 3 min at 4 °C to remove debris and then the supernatant was incubated with anti-HA magnetic beads (Thermo Fisher Scientific, cat. # 88837) for 6 minutes at 20 °C. The bound beads and flowthrough were separated using a magnetic stand. The bound bead fraction was washed 4 times with cold KPBS. The bound bead and flowthrough fractions were both used for LC/MS-based proteomics analyses.

### LC/MS-based proteomic analyses

The bound beads after wash were directly eluted in 100 µl of 5% SDS buffer and trypsin digestion was carried out using S-Trap™ (Protifi, NY) as per manufacturer’s protocol. For the flow-through sample after IP, 100 µl sample was diluted in 5% SDS buffer and trypsin digestion was carried out using S-Trap™. The peptide concentration was measured using the Pierce™ Quantitative Colorimetric Peptide Assay (Thermo Scientific cat. # 23275). The digested peptides were subjected to simple C18 clean up using a C18 disk plug (3M Empore C18) and dried in a speed vac. 1 µg of the peptide was used for LC-MS/MS analysis which was carried out using a nano-LC 1200 system (Thermo Fisher Scientific, San Jose, CA) coupled to Orbitrap Fusion™ Lumos ETD mass spectrometer (Thermo Fisher Scientific, San Jose, CA). The peptides were loaded on a two-column setup using a pre-column trap of 2 cm × 100 µm size (Reprosil-Pur Basic C18 1.9 µm, Dr. Maisch GmbH, Germany) and a 5 cm × 75 µm analytical column (Reprosil-Pur Basic C18 1.9 µm, Dr. Maisch GmbH, Germany) with a 75 min gradient of 5-28% acetonitrile/0.1% formic acid at a flow rate of 750 nl/min. The eluted peptides were directly electro-sprayed into mass spectrometer operated in the data-dependent acquisition (DDA) mode. The full MS scan was acquired in Orbitrap in the range of 300-1400 m/z at 120,000 resolution followed by top 30 MS2 in Ion Trap (AGC 5000, MaxIT 35 ms, HCD 28% collision energy) with 15 sec dynamic exclusion time.

The raw files were searched using Mascot algorithm (Mascot 2.4, Matrix Science) against the *Caenorhabditis elegans* NCBI refseq protein database in the Proteome Discoverer (PD 2.1, Thermo Fisher) interface. The precursor mass tolerance was set to 20 ppm, fragment mass tolerance to 0.5 Da, maximum of two missed cleavage was allowed. Dynamic modification of oxidation on methionine, protein N-terminal Acetylation and deamidation (N/Q) was allowed. The gene product inference and iBAQ-based quantification were carried out using the gpGrouper algorithm (Saltzman et al., 2018). The iBAQ values were normalized to the fraction of total protein iBAQ per experiment defined as ‘iFOT (Fraction of total)’.

### Antibodies

Anti-*C. elegans* LMP-1, HSP-60, CYP-33, SVQ-8 monoclonal antibodies were purchased from Developmental Studies Hybridoma Bank (DHSB). Those antibodies were originally generated by Dr. Michael L. Nonet’s lab (Hadwiger et al., 2010). Anti-β-actin antibody (C4) was purchased from Santa Cruz (sc-47778).

### Microscopy imaging

#### Regular microscopy

Tissue-specific lyso-Tag expression example images (Figure 5A) were captured using Leica DMi8 THUNDER Imaging Systems using 20× objective. Other microscopy images were captured using Olympus FV3000 confocal microscopy system using 60× or 20× objective. *C. elegans* were anesthetized in 1% sodium azide in M9 buffer and placed on 2% agarose pad sandwiched between the glass microscopic slide and coverslip.

#### Fluorescence lifetime microscopy

L1 RNAi sensitive *nre-1(hd20);lin-15b(hd126)* worms were seeded on 3.5cm RNAi plates and raised at 20°C for two days, and then around 20 worms each well were transferred to the 3.5cm RNAi plates containing RNAi bacteria and 0.5 μM of LysoSensor Green DND-189 (Invitrogen™ L7535) and raised for 18h (in dark) at 20°C. The worms were imaged using ISS Q2 Time-resolved Laser Scanning Confocal Nanoscope. The laser excitation wavelength was set at 476nm and the 500-633nm emission filter was used to detect the LysoSenor Green signal. The first pair of intestinal cells of each worm was imaged. The lifetimes of LysoSensor-containing puncta were measured using ISS VistaVision software. The pH of the lysosome was then calculated based on the LysoSensor lifetime-pH calibration curve.

### LysoSensor RNAi screen

The primary screen was performed on 95 lysosomal-enriched candidates shared between the LMP-1 and the CTNS-1 Lyso-IP proteomic profiling datasets. Each RNAi bacteria clone was seeded onto 12-well RNAi plates containing 1 mM IPTG and allowed to dry. The dried plates were then incubated at room temperature overnight to induce dsRNA expression. Synchronized L1 *nre-1(hd20);lin-15b(hd126)* worms were seeded on 12-well RNAi plates and raised at 20 °C for two days, and then around 30 worms each well were transferred to the RNAi plates containing RNAi bacteria and 0.5 μM of LysoSensor Green DND-189. After 18 hours, LysoSensor signals were examined by naked eyes using Nikon SMZ18 fluorescence stereo microscope. The candidates with obvious LysoSensor alteration were selected for the secondary LysosSensor RNAi screen. In the secondary screen, worms stained using LysoSensor Green DND-189 were imaged by the FV3000 confocal laser scanning microscope (At least 15 animals were measured in each condition). The changes in LysoSensor signals in the first pair of intestine cells were quantified by ImageJ (including intensity, size, and number).

### Lysosome distribution quantification

The quantification method was modified from previous publications on lysosomal distribution in mammalian cell lines (Johnson et al., 2016; Willett et al., 2017). Images were first captured using Olympus Fluoview software and imported into Matlab by Bio-Formats tool (Linkert et al., 2010). Next, the cell membrane and nuclear membrane are outlined manually. The algorithm (code included in Supplementary materials) determines the geometric center of the nucleus and radiates at all angles to locate line segments between the nuclear and cell membrane. The line segments are then evenly divided and circled to segment the cytosol into different regions. We calculate: 1. The mean RFP fluorescence distribution across regions (normalized to the mean intensity of the whole cell to avoid variations of RFP expression across cells) (Figure 4D, 4F); 2. The cumulative intensity distribution from the most perinuclear region to the most peripheral region (normalized to the overall intensity of the whole cell) (Supplementary Figure 2B, 2C). The algorithm requires a convex shape of the cell, and most of the gut cells imaged meet this need.

### Lifespan assays

Worms were synchronized by bleach-based egg preparation and subsequent starvation in M9 buffer for over 24 hours. Synchronized L1 worms were placed on the plates, and animals were synchronized again by manual picking at mid L4 stage and marked as Day 0. Approximately 90-120 worms were placed in three parallel plates for each condition. Worms are hence observed and transferred to freshly made RNAi plates every other day. Animals are categorized as alive, death (cessation of movement in response to platinum wire probing) or censored. The statistical analyses were performed with the SPSS23 Kaplan-Meier survival function and the log-rank test. GraphPad Prism 9 was used to graph the results.

### Structural stimulation by AlphaFold2

The structure of R144.6 (UniProt: Q10000) was predicted by AlphaFold2 and downloaded from The AlphaFold Protein Structure Database (https://alphafold.ebi.ac.uk/) (Jumper et al., 2021; Varadi et al., 2022). The molecular graphics of the R144.6 structure was performed with UCSF Chimera (Pettersen et al., 2004).

### Statistical methods

Principal components analysis (PCA) was performed by R package Factextra (Le et al., 2008) with the normalized iFOT abundance of proteins detected as the input. The Pearson correlation matrices and coefficient (r) among replicates were generated by GraphPad Prism 9. The unpaired, two-tailed t-tests for the multiple comparisons were used to calculate the p-values in the correlation analyses by GraphPad Prism 9.

## Data availability

The mass spectrometry data for protein identification have been deposited via the MASSIVE repository (MSV000090909) to the Proteome X change Consortium (http://proteomecentral.proteomexchange.org) with the dataset identifier PXD038865.

Analysis code for Figure 4 D, F and Figure S2 is included in Supplementary code.

## Conflict of Interest

None declared.

## Supporting information

Supplementary Table 1. Lysosome-enriched proteins identified from LMP-1 Lyso-IP using WT worms

Supplementary Table 2. Lysosome-enriched proteins identified from LMP-1 Lyso-IP using lipl-4 Tg worms

Supplementary Table 3. Summary of lifespan analyses

Supplementary Table 4. Lysosome-enriched proteins identified from LMP-1 Lyso-IP using daf-2(lf) mutant

Supplementary Table 5. Lysosome-enriched proteome exhibits tissue-specificity

Supplementary Table 6. Lysosome-enriched proteins identified from CTNS-1 Lyso-IP using WT worms

Supplementary Table 7. LysoSensor screening of lysosome-enriched proteins shared between LMP-1 and the CTNS-1 Lyso-Ips

Supplementary code. Matlab code for lysosome distribution quantification

## ACKNOWLEDGEMENTS

We thank A. Dervisefendic and P. Svay for maintenance support; Mass Spectrometry Proteomics Core of Baylor College of Medicine for mass spectrometry proteomic analysis; Z. Zhou (Baylor College of Medicine) for sharing *unc-76(e911)* strain. Some strains were obtained from the *Caenorhabditis* Genetics Center (CGC), which is funded by NIH Office of Research Infrastructure Programs (P40 OD010440). M.C.W. is currently supported by Howard Hughes Medical Institute, and Y.Y. is currently supported by National Natural Science Foundation of China 32071146.

## AUTHOR CONTRIBUTIONS

Y.Y., S.M.G., Y.G. and M.C.W. conceived the project and designed the experiments. Y.Y., S.M.G., Y.G., P.H., Q. Zhang, J.L., B.J. and S.Y.J. performed experiments. Q. Zhao conducted the structural simulation. D.M.S. and M.A. help design Lyso-IP method in *C. elegans*. S.Y.J. supervised mass spectrometry analysis. Y.Y., S.M.G., Y.G. and M.C.W. wrote the manuscript. Y.Y., S.M.G., Y.G., Q. Zhang, M.A. and M.C.W. edited the manuscript.

## SUPPLEMENTARY FIGURES

**Supplementary Figure 1.**
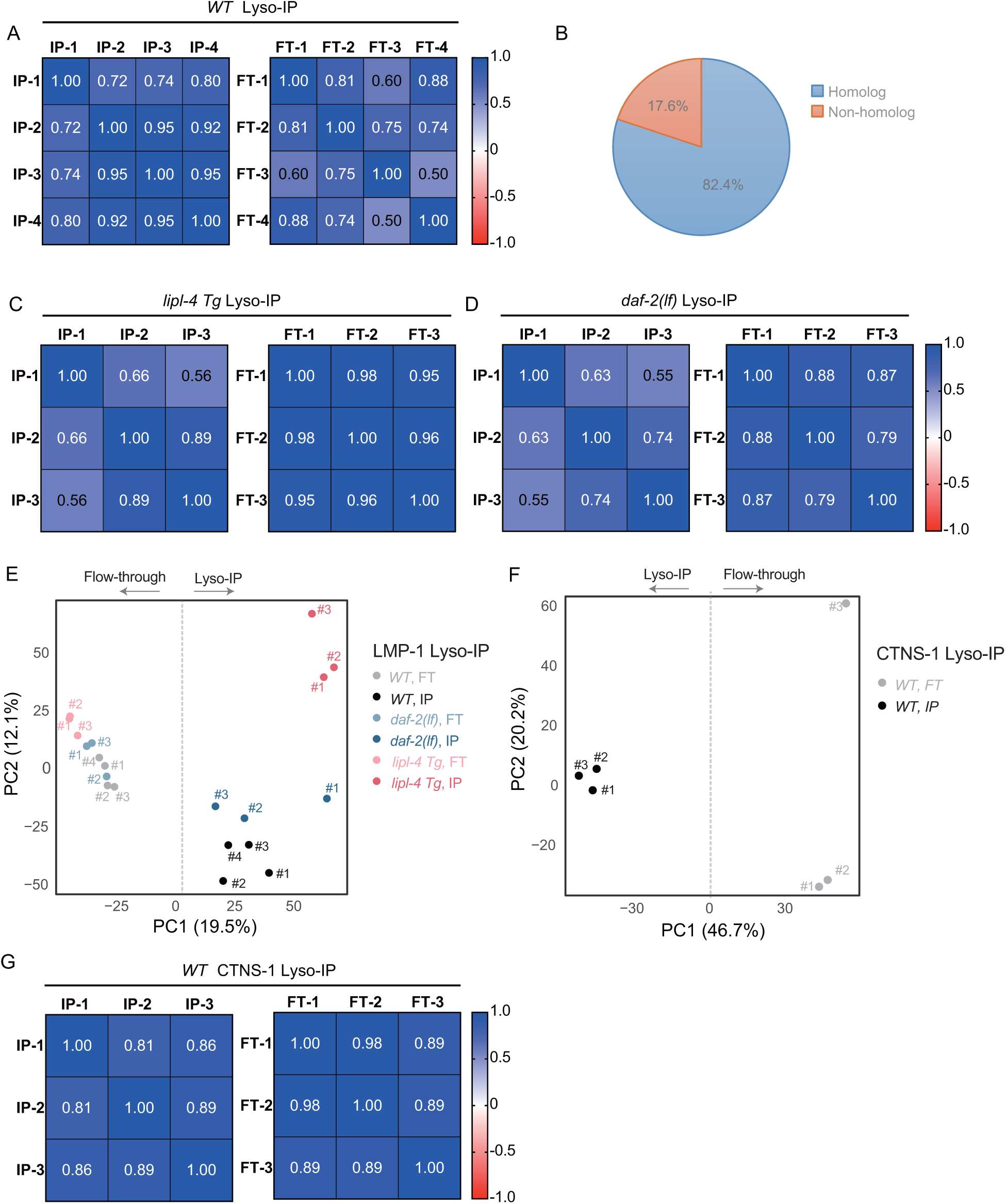
Lyso-IP analyses from worms with different genotypes. (**A**) Correlation analysis of four independent biological replicates of Lyso-IP (IP) and Flow-through (FT) samples from proteomics analyses. **(B)** Pie chart showing the proportion of LMP-1 Lyso-IP candidates from WT worms with mammalian homologs. (**C** and **D)** Correlation analysis of three independent biological replicates of Lyso-IP (IP) and Flow-through (FT) from proteomics analyses of the long-lived *lipl-4* transgenic strain (*lipl-4 Tg*, **C**) and the *daf-2* loss-of-function mutant (*daf-2(lf)*), **D**). **(E)** PCA analysis of Lyso-IP replicates and flow-through controls in LMP-1 Lyso-IP of WT, *lipl-4 Tg* and *daf-2(lf)* worms.| **(F)** PCA analysis of Lyso-IP replicates and flow-through controls of CTNS-1 Lyso-IP of WT worms. **(G)** Correlation analysis of three independent biological replicates of Lyso-IP (IP) and Flow-through (FT) samples from proteomic analyses of CTNS-1 Lyso-IP.

**Supplementary Figure 2.**
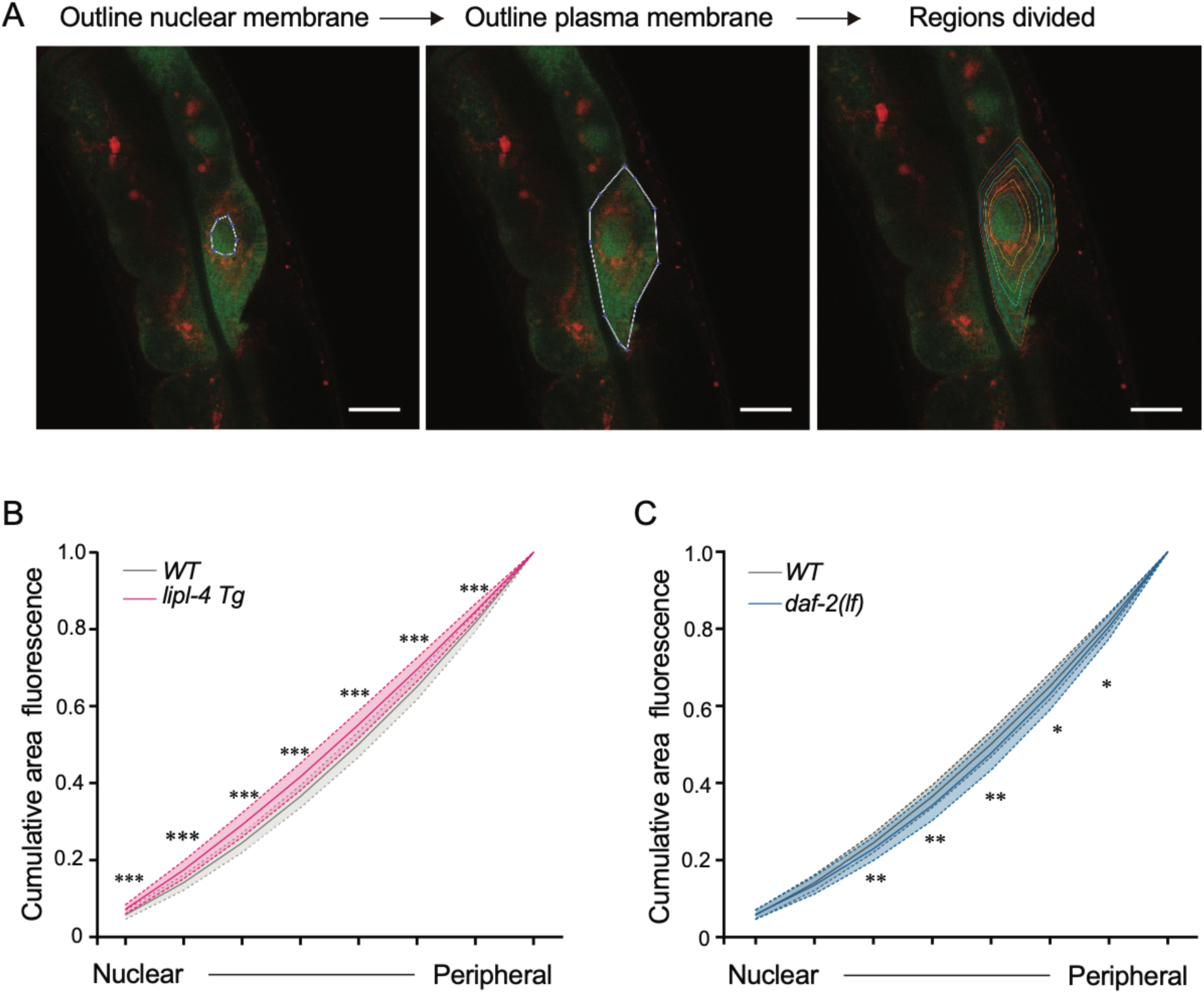
Quantitative analysis of lysosomal positioning. (**A**) Brief summary of the method flow for quantifying the lysosomal distribution in intestinal cells of *C. elegans*. Scale bar=10 μm. **(B, C)** Curve graph showing the normalized accumulated intensity of lysosomal signals from the nuclear to the peripheral region in WT, *lipl-4 Tg* (**B**) and *daf-2(lf)* (**C**) animals. \**p<0.05*; \*\**p<0.01*, \*\*\**p<0.001*, \*\*\*\**p<0.0001*, n.s. *p>0.05* by Student’s t-test (unpaired, two-tailed) for each region. N =50 WT /33 *lipl-4 Tg*, 33 WT/ 28 *daf-2(lf*). Data are represented as mean ± SD. *p* values for (B) (from left to right): 2.65x10^-8^, 3.19 x10^-8^, 7.93 x10^-8^, 3.62 x10^-7^, 4.79 x10^-6^, 2.98 x10^-5^, 4.41 x10^-5^; *p* values for (C) (from left to right): 0.357, 0.0529, 0.00611, 0.00246, 0.00985, 0.0261, 0.0423.

**Supplementary Figure 3.**
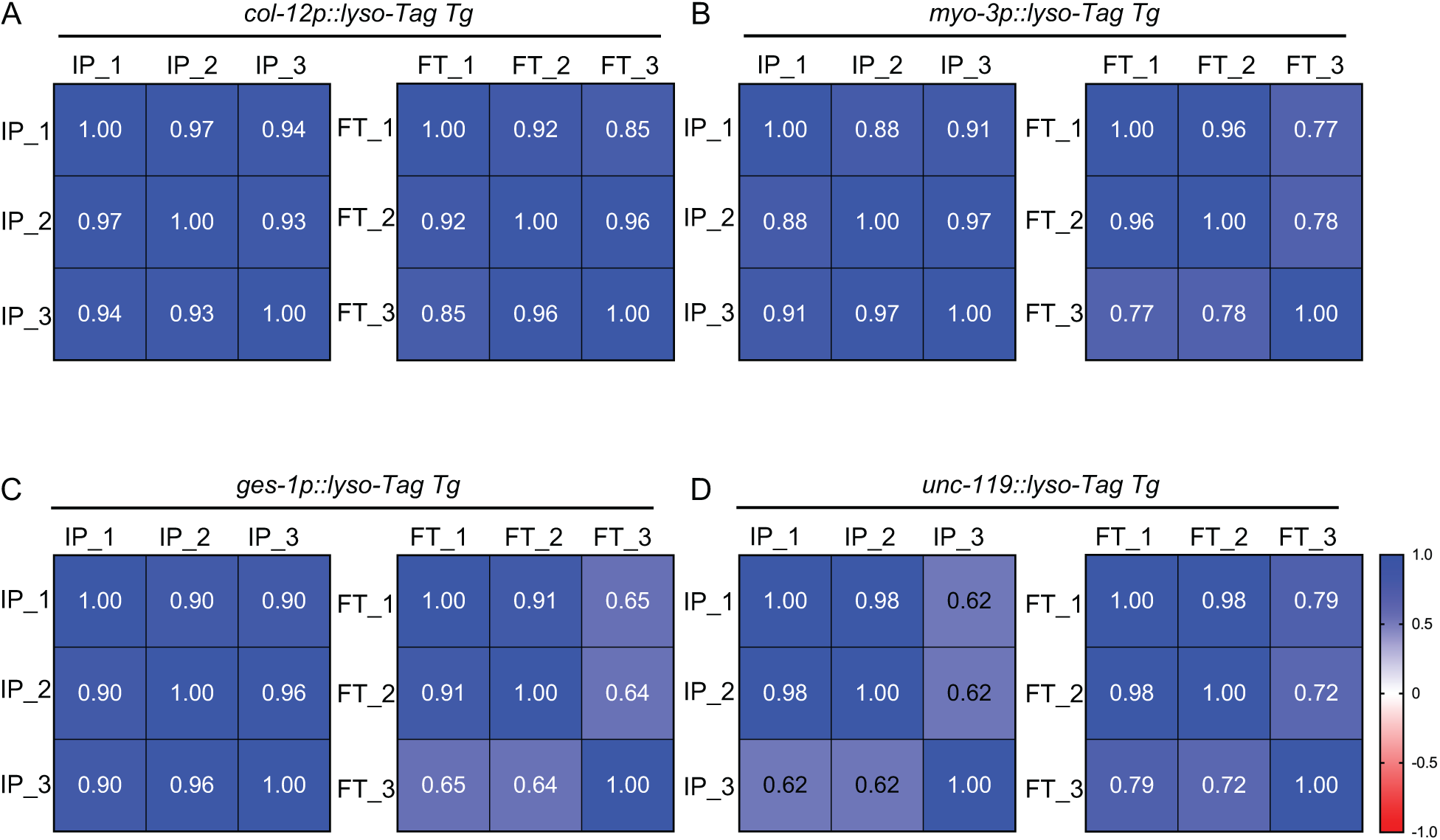
Correlation analyses of tissue-specific Lyso-IPs. (**A-D**) Pearson Correlation matrices of tissue-specific lyso-IP (IP) samples and flow-through (FT) samples show the correlation among three different replicates. **(A)** Hypodermis, **(B)** Muscle, **(C)** Intestine, **(D)** Neuron.

**Supplementary Figure 4.**
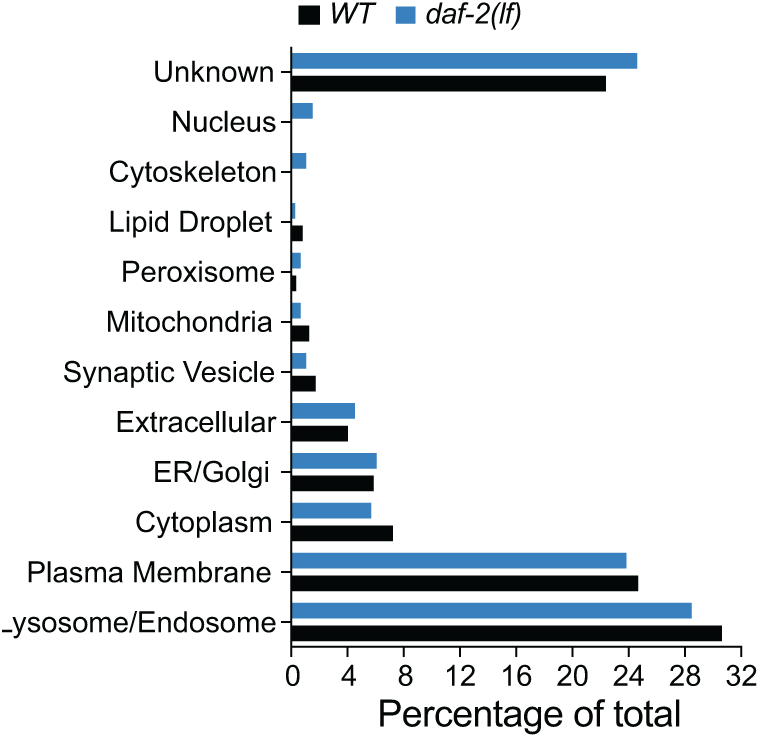
Subcellular category analyses of *daf-2(lf)* candidates. The percentage of proteins with different subcellular localization is compared between lysosome-enriched proteomes from WT and *daf-2(lf)* worms.

**Supplementary Figure 5.**
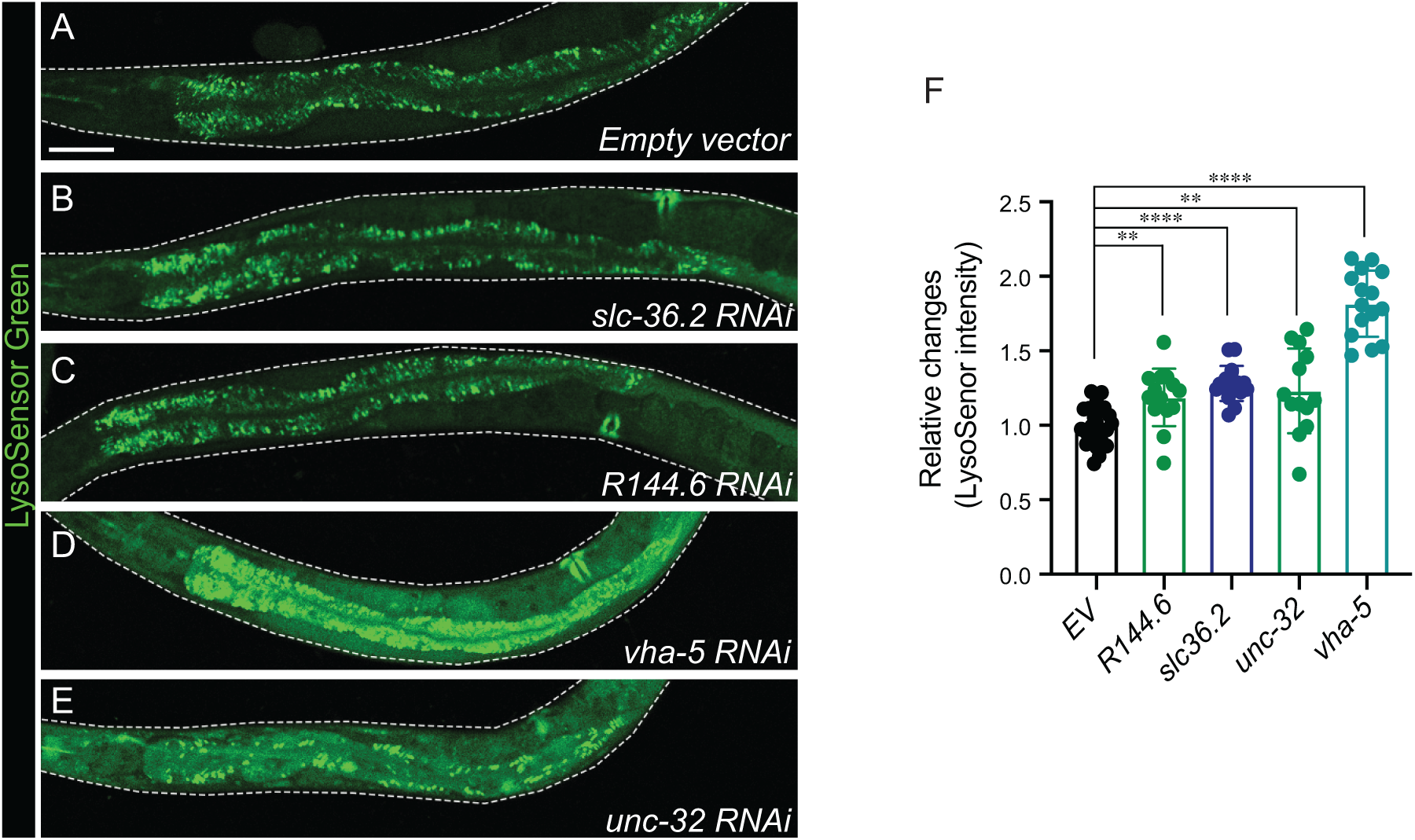
Inactivation of lysosome-enriched candidates increases LysoSensor signal intensity. (**A-E**) Confocal fluorescence microscopy images of intestinal cells in worms stained with LysoSensor DND-189 and treated with *empty vector* (**A**), *slc36.2* RNAi (**B**), *R144.6* RNAi (**C**), *vha-5* RNAi (**D**) and *unc-32* RNAi (**E**). Scale bar=50 μm. **(F)** RNAi knockdown of *slc36.2 (*\*\*\*\**p<0.0001), R144.6 (*\*\**p=0.0011), unc-32 (*\*\**p=0.0019) or vha-5 (*\*\*\*\**p<0.0001)* increases the LysoSensor signal intensity in the area of the first pair of intestinal cells. Data are shown as mean ± SD. Student’s t-test (unpaired, two-tailed) was performed between the *empty vector* group and each RNAi-treated group. n = 24 (*EV*), 14 (*R144.6*), 13 (*slc36.2*), 15 (*unc-32*), 16 (*vha-5*).

**Supplementary Table 1. Lysosome-enriched proteins identified from LMP-1 Lyso-IP using WT worms**

**Supplementary Table 2. Lysosome-enriched proteins identified from LMP-1 Lyso-IP using *lipl-4 Tg* worms**

**Supplementary Table 3. Summary of lifespan analyses**

**Supplementary Table 4. Lysosome-enriched proteins identified from LMP-1 Lyso-IP using *daf-2(lf)* mutant**

**Supplementary Table 5. Lysosome-enriched proteome exhibits tissue-specificity**

**Supplementary Table 6. Lysosome-enriched proteins identified from CTNS-1 Lyso-IP using WT worms**

**Supplementary Table 7. LysoSensor screening of lysosome-enriched proteins shared between LMP-1 and the CTNS-1 Lyso-Ips**

**Supplementary code. Matlab code for lysosome distribution quantification**

## Notes

### Competing Interest Statement

The authors have declared no competing interest.

### Summary of Updates

Data availability statement provided. Some minor revisions on manuscript.

