## Supplementary code. Matlab code for lysosome distribution quantification for "Organelle proteomic profiling reveals lysosomal heterogeneity in association with longevity": Supplementary code_lysosome quantification.m

```
addpath '#path to image directory'
clear;
[file,path]=uigetfile('*.oir','select a oir file'),
filename=fullfile(path,file);
datain = bfopen(filename);
datafigure1=datain{1,1};
gfp=datafigure1{1,1};
rfp=datafigure1{2,1};

figure;
C = imfuse(gfp,rfp,'ColorChannels',[2 1 0]);
imshow(C,[]);

#display the image
#get the outline for nucleus and cell membrane
i = 1;
nn = 1; 
 while(i<=nn)
       keydoprown = waitforbuttonpress;
       if (keydoprown == 1) 
              [bw1,xi1,yi1]=roipoly;
             i = i+1;
       end
 end    
   

 
i = 1;
 while(i<=nn)
       keydoprown = waitforbuttonpress;
       if (keydoprown == 1) 
              [bw2,xi2,yi2]=roipoly;
               close all;
             i = i+1;
       end
 end    
 

picsize=size(gfp);

polyin=polyshape(xi1,yi1);
[x,y]=centroid(polyin);

xf=zeros(360,9);
yf=zeros(360,9);

for n = 1:89
xline=[x x-tand(90-n)*y];
yline=[y 0];
[xt1,yt1] = polyxpoly(xline,yline,xi1,yi1);
[xt2,yt2] = polyxpoly(xline,yline,xi2,yi2);
xf(n,:)=linspace(xt1,xt2,9);
yf(n,:)=linspace(yt1,yt2,9);
end


xline6=[x x];
yline6=[y 0];
[x61,y61] = polyxpoly(xline6,yline6,xi1,yi1);
[x62,y62] = polyxpoly(xline6,yline6,xi2,yi2);
xf(90,:)=linspace(x61,x62,9);
yf(90,:)=linspace(y61,y62,9);

for n = 1:89
xline=[x x+tand(n)*y];
yline=[y 0];
[xt1,yt1] = polyxpoly(xline,yline,xi1,yi1);
[xt2,yt2] = polyxpoly(xline,yline,xi2,yi2);
xf(n+90,:)=linspace(xt1,xt2,9);
yf(n+90,:)=linspace(yt1,yt2,9);
end


xline12=[x picsize(1)];
yline12=[y y];
[x121,y121] = polyxpoly(xline12,yline12,xi1,yi1);
[x122,y122] = polyxpoly(xline12,yline12,xi2,yi2);
xf(180,:)=linspace(x121,x122,9);
yf(180,:)=linspace(y121,y122,9);

for n = 1:89
xline=[x x+tand(90-n)*(picsize(1)-y)];
yline=[y picsize(1)];
[xt1,yt1] = polyxpoly(xline,yline,xi1,yi1);
[xt2,yt2] = polyxpoly(xline,yline,xi2,yi2);
xf(n+180,:)=linspace(xt1,xt2,9);
yf(n+180,:)=linspace(yt1,yt2,9);
end


xline18=[x x];
yline18=[y picsize(1)];
[x181,y181] = polyxpoly(xline18,yline18,xi1,yi1);
[x182,y182] = polyxpoly(xline18,yline18,xi2,yi2);
xf(270,:)=linspace(x181,x182,9);
yf(270,:)=linspace(y181,y182,9);

for n = 1:89
xline=[x x-tand(n)*(picsize(1)-y)];
yline=[y picsize(1)];
[xt1,yt1] = polyxpoly(xline,yline,xi1,yi1);
[xt2,yt2] = polyxpoly(xline,yline,xi2,yi2);
xf(n+270,:)=linspace(xt1,xt2,9);
yf(n+270,:)=linspace(yt1,yt2,9);
end


xline24=[x 0];
yline24=[y y];
[x241,y241] = polyxpoly(xline24,yline24,xi1,yi1);
[x242,y242] = polyxpoly(xline24,yline24,xi2,yi2);
xf(360,:)=linspace(x241,x242,9);
yf(360,:)=linspace(y241,y242,9);


%imshow(bw4,[]);
%hold on;
%hold off;


 %imshow(bw4,[]);
%hold on;
%imshow(rfp,[]);
%hold on;

%plot(xi1,yi1,'g');
%plot(xf(:,2),yf(:,2),'g');
%plot(xf(:,3),yf(:,3),'r');
%plot(xf(:,9),yf(:,5),'b');

hold off;

bwcal1 = poly2mask(xf(:,1),yf(:,1),picsize(1),picsize(2));
bwcal2 = poly2mask(xf(:,2),yf(:,2),picsize(1),picsize(2));
bwcal3 = poly2mask(xf(:,3),yf(:,3),picsize(1),picsize(2));
bwcal4 = poly2mask(xf(:,4),yf(:,4),picsize(1),picsize(2));
bwcal5 = poly2mask(xf(:,5),yf(:,5),picsize(1),picsize(2));
bwcal6 = poly2mask(xf(:,6),yf(:,6),picsize(1),picsize(2));
bwcal7 = poly2mask(xf(:,7),yf(:,7),picsize(1),picsize(2));
bwcal8 = poly2mask(xf(:,8),yf(:,8),picsize(1),picsize(2));
bwcal9 = poly2mask(xf(:,9),yf(:,9),picsize(1),picsize(2));

region1=bwcal2&(~bwcal1);
region2=bwcal3&(~bwcal2);
region3=bwcal4&(~bwcal3);
region4=bwcal5&(~bwcal4);
region5=bwcal6&(~bwcal5);
region6=bwcal7&(~bwcal6);
region7=bwcal8&(~bwcal7);
region8=bwcal9&(~bwcal8);


pin1=rfp(region1);
pin2=rfp(region2);
pin3=rfp(region3);
pin4=rfp(region4);
pin5=rfp(region5);
pin6=rfp(region6);
pin7=rfp(region7);
pin8=rfp(region8);


sum5=[sum(pin1),sum(pin2),sum(pin3),sum(pin4),sum(pin5),sum(pin6),sum(pin7),sum(pin8)];

sum4=[mean(pin1),mean(pin2),mean(pin3),mean(pin4),mean(pin5),mean(pin6),mean(pin7),mean(pin8)];

for n= 1: 8
   eval(['max',num2str(n),'=size(pin',num2str(n),');']); 
end

sumsize=[max1(1),max2(1),max3(1),max4(1),max5(1),max6(1),max7(1),max8(1)];
writecell({file,sum4,sumsize,sum5},'quantification.xls','WriteMode','append');
#save nuclear and cell membrane location if needed for other anaylsis

# bb=[xi1,yi1];
# bb2=[xi2,yi2];
# savename=[file 'in.xls'];
#writematrix(bb,savename);
#savename=[file 'out.xls'];
#writematrix(bb2,savename);
```
